# Transcriptional Landscape of Ectomycorrhizal Fungi and Their Host Provide Insight into N Uptake from Forest Soil

**DOI:** 10.1101/2021.07.23.453179

**Authors:** Carmen Alicia Rivera Pérez, Dennis Janz, Dominik Schneider, Rolf Daniel, Andrea Polle

## Abstract

Mineral nitrogen (N) is a major nutrient showing strong fluctuations in the environment due to anthropogenic activities. Acquisition and translocation of N to forest trees is achieved by highly diverse ectomycorrhizal fungi (EMF) living in symbioses with their host roots. Here, we examined colonized root tips to characterize the entire root-associated fungal community by DNA metabarcoding-Illumina sequencing of the fungal ITS2 molecular marker and used RNA sequencing to target metabolically active fungi and the plant transcriptome after N application. The study was conducted with beech (*Fagus sylvatica* L), a dominant tree species in central Europe, grown in native forest soil. We demonstrate strong enrichment of ^15^N from nitrate or ammonium in the ectomycorrhizal roots by stable isotope labeling. The relative abundance of the EMF members in the fungal community was correlated with their transcriptional abundances. The fungal metatranscriptome covered KEGG and KOG categories similar to model fungi and did not reveal significant changes related to N metabolization but species-specific transcription patterns, supporting trait stability. In contrast to the resistance of the fungal metatranscriptome, the transcriptome of the host exhibited dedicated nitrate- or ammonium-responsive changes with upregulation of transporters and enzymes required for nitrate reduction and drastic enhancement of glutamine synthetase transcript levels, indicating channeling of ammonium into the pathway for plant protein biosynthesis. Our results support that self-composed fungal communities associated with tree roots buffer nutritional signals in their own metabolism but do not shield plants from high environmental N.

**IMPORTANCE:** Although EMF are well known for their role in supporting tree N nutrition, the molecular mechanisms underlying N flux from the soil solution into the host through the ectomycorrhizal pathway remain widely unknown. Furthermore, ammonium and nitrate availability in the soil solution is subject to constant oscillations that create a dynamic environment for the tree roots and associated microbes during N acquisition. Therefore, it is important to understand how root-associated mycobiomes and the tree roots handle these fluctuations. We studied the response of the symbiotic partners by screening their transcriptomes after a sudden environmental flux of nitrate or ammonium. We show that the fungi and the host respond asynchronously, with the fungi displaying resistance to increased nitrate or ammonium, and the host dynamically metabolizing the supplied N sources. This study provides insights into the molecular mechanisms of the symbiotic partners operating under N enrichment in a multidimensional symbiotic system.

---

**S**oil N availability is generally a main limiting factor for primary productivity across terrestrial ecosystems including temperate forests (1, 2). In forest soil, soluble mineral N pools consist of nitrate and ammonium, whose quantities fluctuate in time and space, depending on soil properties, meteorological conditions, anthropogenic N inputs and biological processes such as mineralization, immobilization, and denitrification (3–12). While nitrate ions are highly mobile in soil solution and easily lost by leaching, ammonium cations are generally bound to soil colloids and retained in topsoil (13, 14). Consequently, mineral N nutrition of plants and microbes must cope with dynamic N availabilities in the environment.

The mutualistic association of certain soil EMF species with the root tips of forest trees is an ecological advantage to support nutrition of the host from variable environmental N sources (15–20). The vast majority of the root systems of individual trees in temperate forests are naturally colonized by a diverse spectrum of EMF species forming compound organs known as ectomycorrhizas and variably composed fungal communities (21–24). These ectomycorrhizas consist of root and fungal cells that mediate bidirectional nutrient exchange. EMF acquire N from the environment and transfer it to the root and receive host-derived carbon in return (25, 26). In self-assembled ectomycorrhizas, EMF show strong interspecific differences for N acquisition (27, 28). Early laboratory experiments showed that when the mycelium of EMF colonizing the roots of *Pinus sylvestris* and *Fagus sylvatica* was supplied with either ammonium or nitrate, the N sources became predominantly incorporated into the amino acids glutamate, glutamine, aspartate, asparagine and alanine (29, 30). When ammonium and nitrate were supplied at equimolar quantities to the mycelium of *Paxillus involutus*, ammonium incorporation into amino acids occurred in the fungus and nitrate remained almost unchanged, suggesting that EMF assimilate ammonium more readily than nitrate into amino acids prior to delivering it to the plant (31). Despite known discrimination between nitrate and ammonium (32, 33), most EMF have a widespread ability to metabolize nitrate (34, 35). Silencing the nitrate reductase (NR) gene in *Laccaria bicolor* impaired the formation of mycorrhizas with poplar (36) implying an important role of EMF in nitrate acquisition for the host.

The process of N transfer to the host through the mycorrhizal pathway starts at the soil-fungal interface, where different N forms are taken up from the soil solution by fungal membrane transporters, N is then translocated through the fungal mantle, which wrapps the root tip, into the intraradical hyphae, and finally exported to the symbiotic interface becoming available for the plant (37–42). Studies on *Amanita muscaria, Hebeloma cylindrosporum, L. bicolor* and *Tuber melanosporum* have led to the hypotheses that ammonium is exported from the intraradical hyphae to the symbiotic interface through Ammonia/Ammonium Transport Out (Ato) proteins, voltage-dependent cation channels, and aquaporins (37, 43–46), and that amino acid export could occur through Acids Quinidine Resistance 1 proteins in *L. bicolor* and *H. cylindrosporum* (38, 44, 47). Moreover, the EMF-mediated import of ammonium and nitrate into the roots is supported by upregulation of ammonium transporters (43) and nitrate transporter (*NRT*) genes in ectomycorrhizal poplar roots like *PttNRT2*.*4A* with *A. muscaria* (48) and *PcNRT1*.*1 and PcNRT2*.*1* with *P. involutus* (49).

Once nitrate is taken up by NRTs, it is intracellularly reduced to nitrite by NR, then to ammonium by NiR and ammonium is ultimately incorporated into glutamine and glutamate (47, 50, 51). In the cyclic GS-GOGAT pathway, glutamine synthetase (GS) catalyzes the formation of glutamine by transfer of ammonium to glutamate. Then glutamate synthase (GOGAT) transfers the amino group from glutamine to 2-oxoglutarate generating two molecules of glutamate, whereas in the alternative pathway, the enzyme glutamate dehydrogenase (GDH) catalyzes the reductive amination of one molecule of 2-oxoglutarate using ammonium to generate one molecule of glutamate (50, 51). Both GS/GOGAT and GDH pathways operate in EMF but variations are common among species or symbiotic systems depending on the plant and fungal partners (52–54). In contrast to EMF, in plants the GS/GOGAT pathway predominates and GDH plays a minor role in ammonium incorporation into organic N forms (55). Currently, the molecular processes used by EMF for supplying mineral N to the host in field conditions are unknown. Uncovering these molecular activities will enable a better understanding of tree N nutrition and N cycling in the ecosystem.

Despite the well-recognized importance of the mycorrhizal pathway as a relevant route whereby tree roots acquire N, knowledge of the molecular mechanisms operating in the uptake, transport, and delivery of N to the host is limited to a few model EMF. It is also unknown how EMF and the colonized root cells respond to variation in mineral N availabilities. The “1000 Fungal Genomes Project” (56) along with the *Fagus sylvatica* genome (57) provide a platform for disentangling fungal and plant transcriptional profiles in self-assembled communities engaged in active symbioses. We took advantage of new tools to unravel these responses in natural forest soil administering a N dose corresponding to 29 kg N ha^-1^ yr^-1^, a quantity in the range of an N saturated beech forest (58, 59). To control N uptake and to distinguish responses to different N forms, we fertilized with either ^15^N-labeled ammonium or ^15^N-labeled nitrate and then studied transcriptional responses separately for EMF and the host trees using ectomycorrhizal root tips (EMRTs). We used DNA-barcoding to describe the composition of the root-associated fungal community and RNA sequencing to capture the metabolically active fungi. We hypothesized that (i) the fungal community structure is unaffected after short-term exposure to elevated N and that (ii) the transcriptional responses of metabolically active EMF reveal molecular activities related to uptake and assimilation of nitrate and ammonium. Since nitrate assimilation requires a series of reduction steps into ammonium before its incorporation into amino acids, both distinct and overlapping responses to nitrate and ammonium availability were expected to be imprinted in the transcription profiles of the symbiotic partners. Furthermore, we hypothesized that (iii) EMF buffer environmental fluctuations in N for the plant resulting in strong N-induced responses in the fungal metatranscriptome but only marginal effects in the root transcriptome, or alternatively that (iv) the entire symbiotic system forms a “holobiont” where the host and the EMF partners display synchronized and similar N-responses.

## RESULTS

### Abundance of root-associated fungal genera corresponds to transcriptional abundance

The global fungal community associated with beech roots in this experiment was dominated by six genera containing ectomycorrhizal (*Amanita:* 7.18%, *Cenococcum:* 9.05%, *Scleroderma:* 4.83%, *Xerocomus:* 29.17%), ericoid (*Oidiodendron:* 1.09%) and saprotrophic fungi (*Mycena:* 3.75%) (Fig. 1A; Data set 1). The remaining taxa were rare (< 1% per genus) and belonged to the phyla of Ascomycota (2.31%), Basidiomycota (2.51%), Mucoromycota (0.11%), Mortierellomycota (0.02%), and fungi of unknown phylogenetic lineage (39.98%) (Fig. 1A; Data set 1). We did not detect any significant effects of short-term ammonium or nitrate treatment on fungal OTU richness (F^2,9^= 0.288, p= 0.756), on Shannon diversity (F^2,9^= 0.437, p= 0.659) (Table S1), or on the composition of the fungal OTU assemblages (R^2^= 0.146, pseudo-F^2,9^= 0.767, p= 0.861, permutations = 9999, adonis, Fig. S1A).

**FIG 1.**
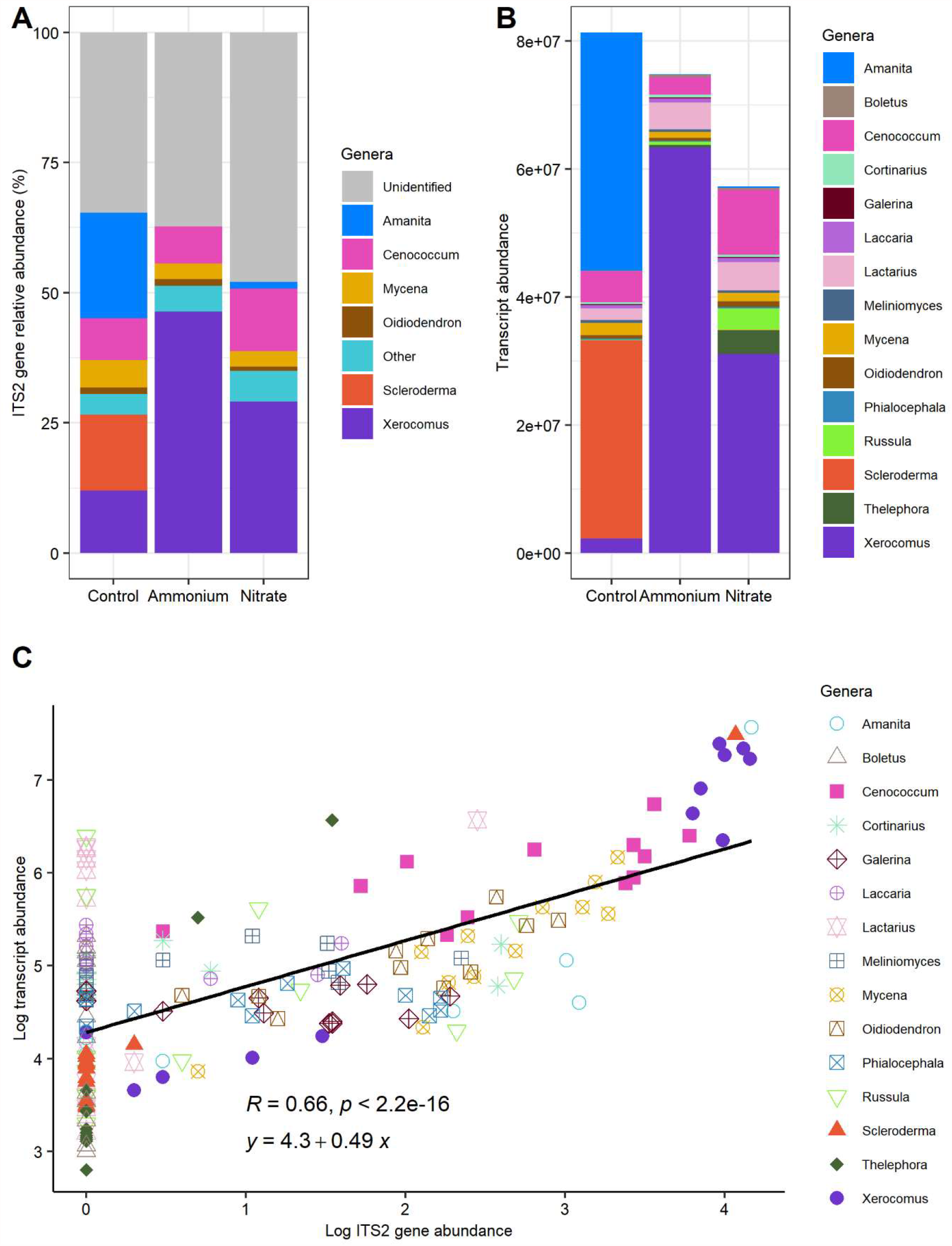
Relative abundance of root-associated (RAF) fungi based on ITS2 barcoding (A), raw counts of the metabolically active fungi based on RNA sequencing characterized by taxonomy (B) and Pearson correlation between DNA-based and RNA-based abundances of the fungal genera (C). RAF were studied on roots of European beech (*Fagus sylvatica*) grown in native forest soil, treated with either water (control), ammonium or nitrate for two days before harvest, (n = 4 per treatment).

Similarly as for the fungal OTUs, we aggregated the RNA counts of ectomycorrhizal fungi belonging the same genus (Fig. 1B). The transcript abundances obtained for individual genera were variable within replicates and treatment groups. However, there were no significant differences among the fungal metatranscriptomes in the nitrate, ammonium or control treatments (R^2^= 0.198, pseudo-F_2,9_= 1.110, p= 0.353, permutations = 9999, adonis) (Fig. 1B; Fig. S1B). The transcript abundance of a specific fungal genus was strongly correlated with the ITS-based abundance of that same genus (R= 0.66, p< 0.001, Pearson, Fig. 1C), supporting that the metabolic activities of abundant fungi associated with the beech roots were reflected. Fungi with low abundances as determined by the DNA-based approach also showed significant transcript abundances (Fig. 1C), implying that low-abundant fungi still may contribute significantly to the molecular activities of the root mycobiome.

### Fungal metatranscriptomes cover fungal metabolism which hardly respond to N treatments

The RNA data containing ectomycorrhizal, ericoid mycorrhiza, endophyte and saprotrophic fungi comprised a total of 175,531 transcript identifiers or gene models, covering 3,759 unique Eukaryotic Orthologous Groups of protein identifiers (KOGs). From these, 122,437 transcript ids (covering 3,708 unique KOGs) belong purely to the EMF (Data set 2). After aggregating the fungi by KOGs into a metatranscriptome and normalizing in DESeq2, the full list fungal metatranscriptome (17 fungi, see Table 1) resulted in 3,619 unique KOGs, whereas the EMF-specific metatranscriptome (13 EMF species, see Table 1) comprised 3,593 KOGs (Data set 3). We evaluated the molecular functions of the EMF metatranscriptome according to KOG functional classifications. All 25 KOG functions were represented and categorized into “cellular processing and signaling” (1,159 KOGs), “information, storage and processing” (956 KOGs), “Metabolism” (796 KOGs), “poorly characterized” (817 KOGs), and multiple function assignment (135 KOGs) (Fig. 2). The frequencies of these functional classifications roughly reflected the same pattern of KOG frequencies present *in silico* in the model EMF *L. bicolor* and that of *Laccaria* sp. on the beech roots (Fig. 2).

**TABLE 1.**
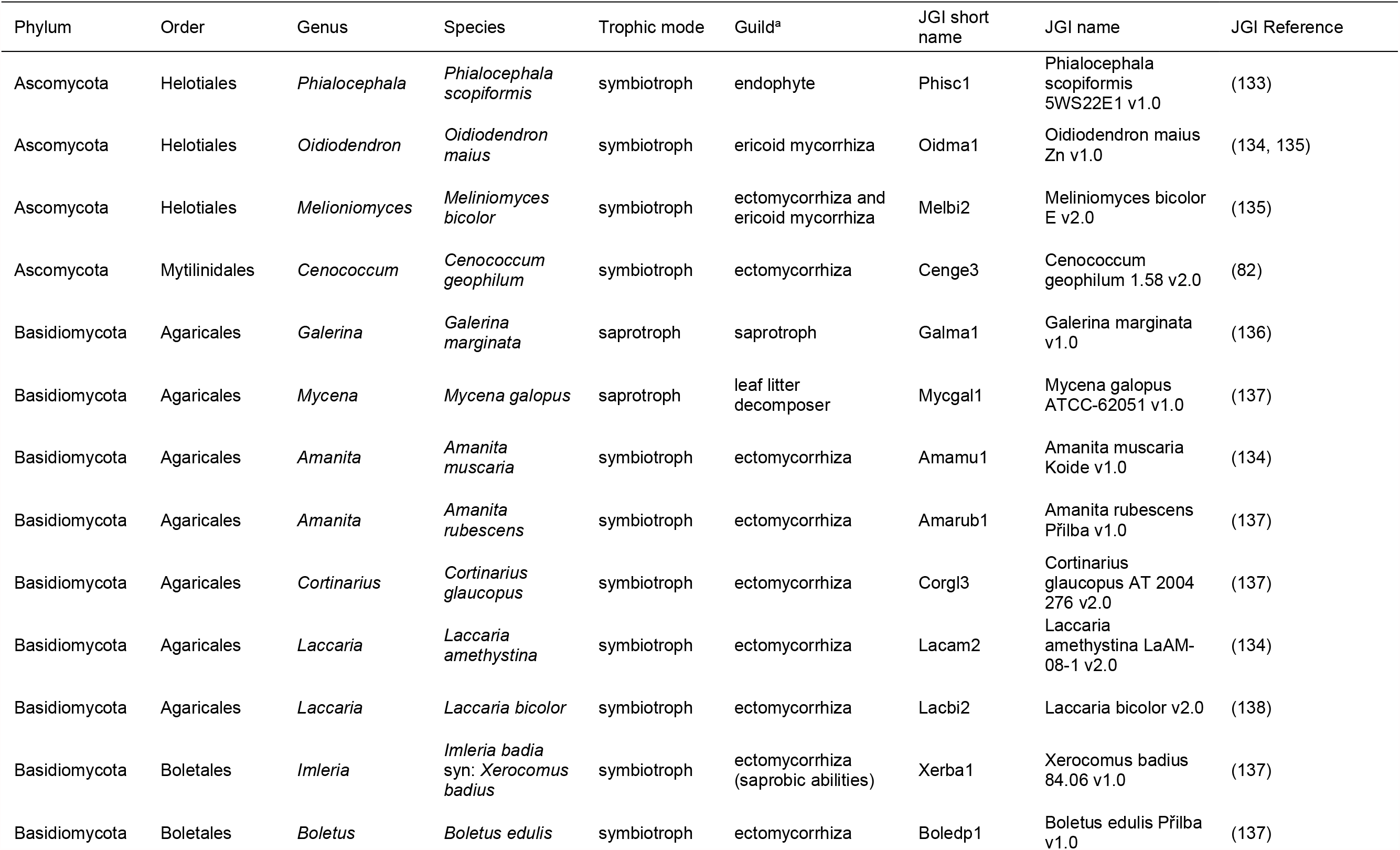
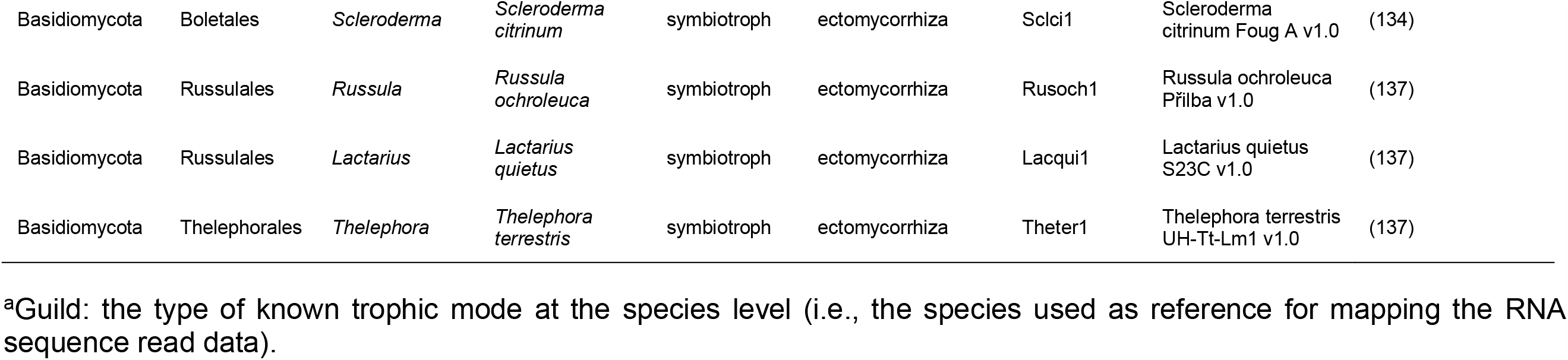
Taxonomy of the genera representing the beech root-associated fungal community and the reference species chosen from the JGI MycoCosm database for mapping the RNA sequencing data

**FIG 2.**
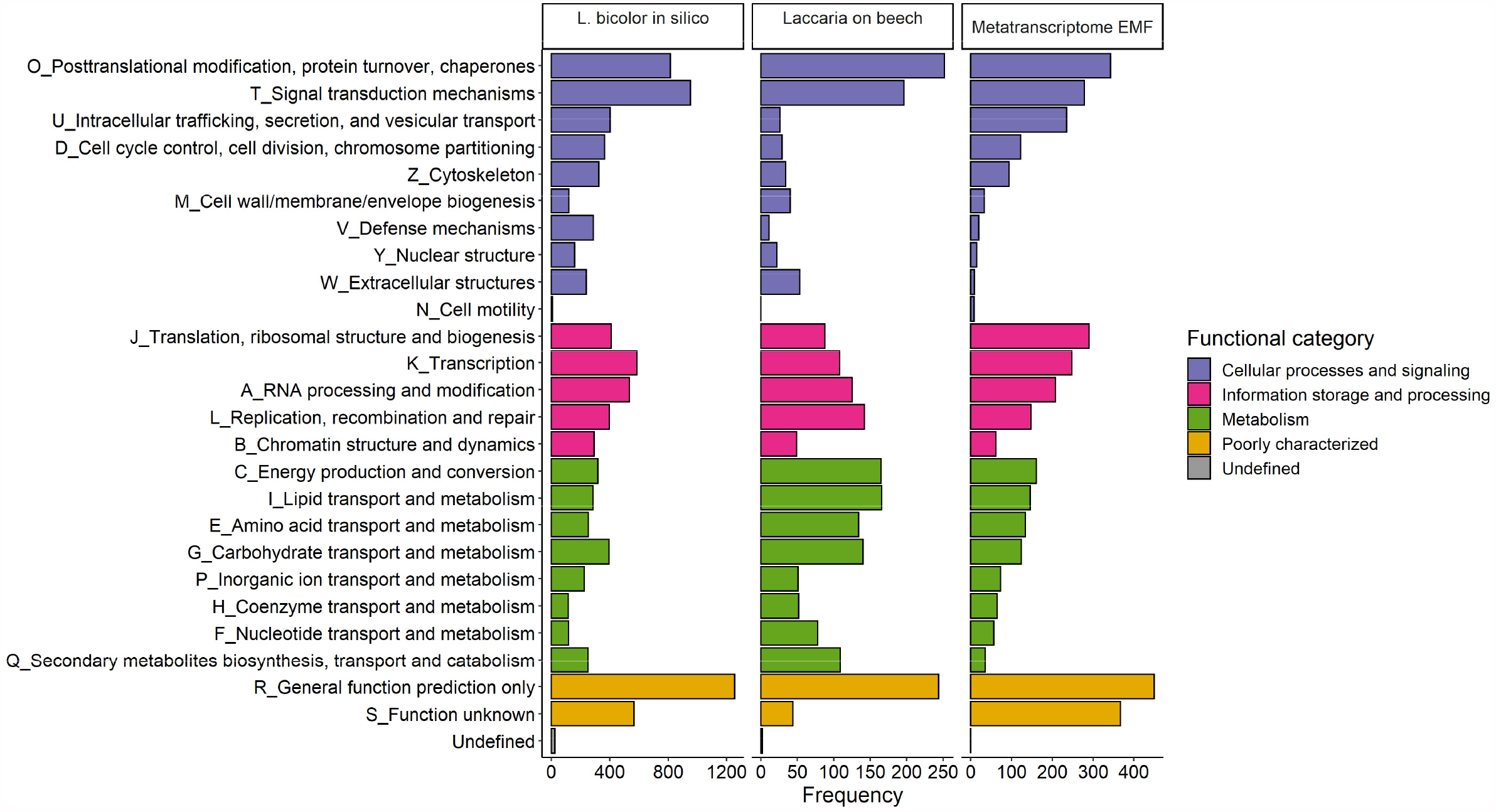
Functional classification of the ectomycorrhizal fungi (EMF) metatranscriptome according to KOG functional groups. The figure shows the distribution of KOG functions for the model ectomycorrhizal fungus *Laccaria bicolor* (*in silico* analyses of the published genome (138)), KOG functions in the transcriptome of the genus *Laccaria* in this experiment (*Laccaria* on beech), and in the entire ectomycorrhizal fungal metatranscriptome in this experiment (Metatranscriptome EMF).

We further tested with DESeq2 whether the KOGs belonging to the full list fungal metatranscriptome or only to the EMF metatranscriptome were significantly differentially expressed in response to ammonium or nitrate treatment relative to the controls. In response to ammonium, not a single KOG was significantly affected (Data set 3). In response to nitrate, one differentially expressed KOG was detected (KOG4381) in the full list fungal metatranscriptome and two KOGs (KOG4381 and KOG4431) in the EMF metatranscriptome (Data set 3). KOG4381 (RUN domain-containing protein) was 578-fold (p _adjusted_ = 0.023943) and 550-fold (p_adjusted_ = 0.020832) upregulated in the full list and in the EMF metatranscriptomes, respectively (Data set 3). This KOG’s function is “signal transduction mechanisms” under the “cellular processes and signaling” category. KOG4431 (uncharacterized protein induced by hypoxia) was 2.27-fold decreased (p_ajusted_ = 0.020832) in response to nitrate and has “poorly characterized function” (Data set 3). Two fungi (*Cenococcum geophilum* and *Xerocomus badius*) occurred in all samples (Fig. 1B), but because of overall low transcriptome coverage, we did not test differential responses to N treatments for specific fungi.

Mapping the EMF metatranscriptome to the KEGG pathway database with *L. bicolor* as reference revealed 108 metabolic pathways, including “biosynthesis of amino acids,” “carbon metabolism,” and “nitrogen metabolism” (Table S2). From a total of 952 unique EC numbers, the complete (866) were mapped and the partial (86) were excluded to avoid inaccurate multiple reaction assignments (60). KEGG pathway enrichment analysis pooling all treatments revealed putative metabolic functions of the EMF metatranscriptome with eleven significantly enriched pathways (FDR P_adjusted_ <0.05), mainly for energy, carbon, amino acid and N metabolism: “glycolysis/glucogenesis,” “pentose phosphate pathway,” “pyruvate metabolism,” “amino sugar and nucleotide sugar metabolism,” “pyrimidine metabolism,” “biosynthesis of amino acids,” “arginine biosynthesis” (Table 2). While “nitrogen metabolism” was covered but not significant (P = 0.063) with the enzymes GS (EC 6.3.1.2), GDH (EC 1.4.1.2), nitrilase (EC 3.5.5.1), and carbonic anhydrase (EC 4.2.1.1). After manually searching the complete fungal transcriptional database (Data set 2), transcripts encoding proteins and enzymes for fungal N uptake and metabolism were discovered. These clustered according to the fungal species instead of putative transporter/enzyme function (Fig. 3). The samples did not clearly cluster according to treatments but formed two main clusters, one containing the majority of nitrate- and ammonium-treated samples (6/8), the controls (4/4), and 2 N-treated samples. However, these differences were not significant (R^2^= 0.176, pseudo-F_2,9_= 0.96161, p = 0.475, adonis).

**TABLE 2.** KEGG pathway enrichment analysis of the ectomycorrhizal fungi metatranscriptome^b^

| Term | Term name | P adjusted |
| --- | --- | --- |
| KEGG:01100 | Metabolic pathways | 5.3E-17 |
| KEGG:01110 | Biosynthesis of secondary metabolites | 7.3E-11 |
| KEGG:00230 | Purine metabolism | 8.1E-07 |
| KEGG:01230 | Biosynthesis of amino acids | 9.4E-05 |
| KEGG:01200 | Carbon metabolism | 1.3E-04 |
| KEGG:00010 | Glycolysis / Gluconeogenesis | 2.5E-03 |
| KEGG:00620 | Pyruvate metabolism | 5.9E-03 |
| KEGG:00520 | Amino sugar and nucleotide sugar metabolism | 2.4E-02 |
| KEGG:00030 | Pentose phosphate pathway | 2.5E-02 |
| KEGG:00220 | Arginine biosynthesis | 2.9E-02 |
| KEGG:00240 | Pyrimidine metabolism | 2.9E-02 |
| KEGG:00680 | Methane metabolism | 6.3E-02 |
| KEGG:00261 | Monobactam biosynthesis | 6.3E-02 |
| KEGG:00250 | Alanine, aspartate and glutamate metabolism | 6.3E-02 |
| KEGG:00910 | Nitrogen metabolism | 6.3E-02 |

<sup>b</sup>Enrichment analysis was performed in g:Profiler against the ascomycete *Aspergillus oryzae* (version e100\_eg47\_p14\_7733820; date: 10/20/2020) since the model organism *Laccaria bicolor* is not available in g:Profiler. Terms indicate the KEGG pathways to which EC numbers are mapped; Term name indicates the KEGG pathways; P adjusted is the FDR corrected p value.

**FIG 3:**
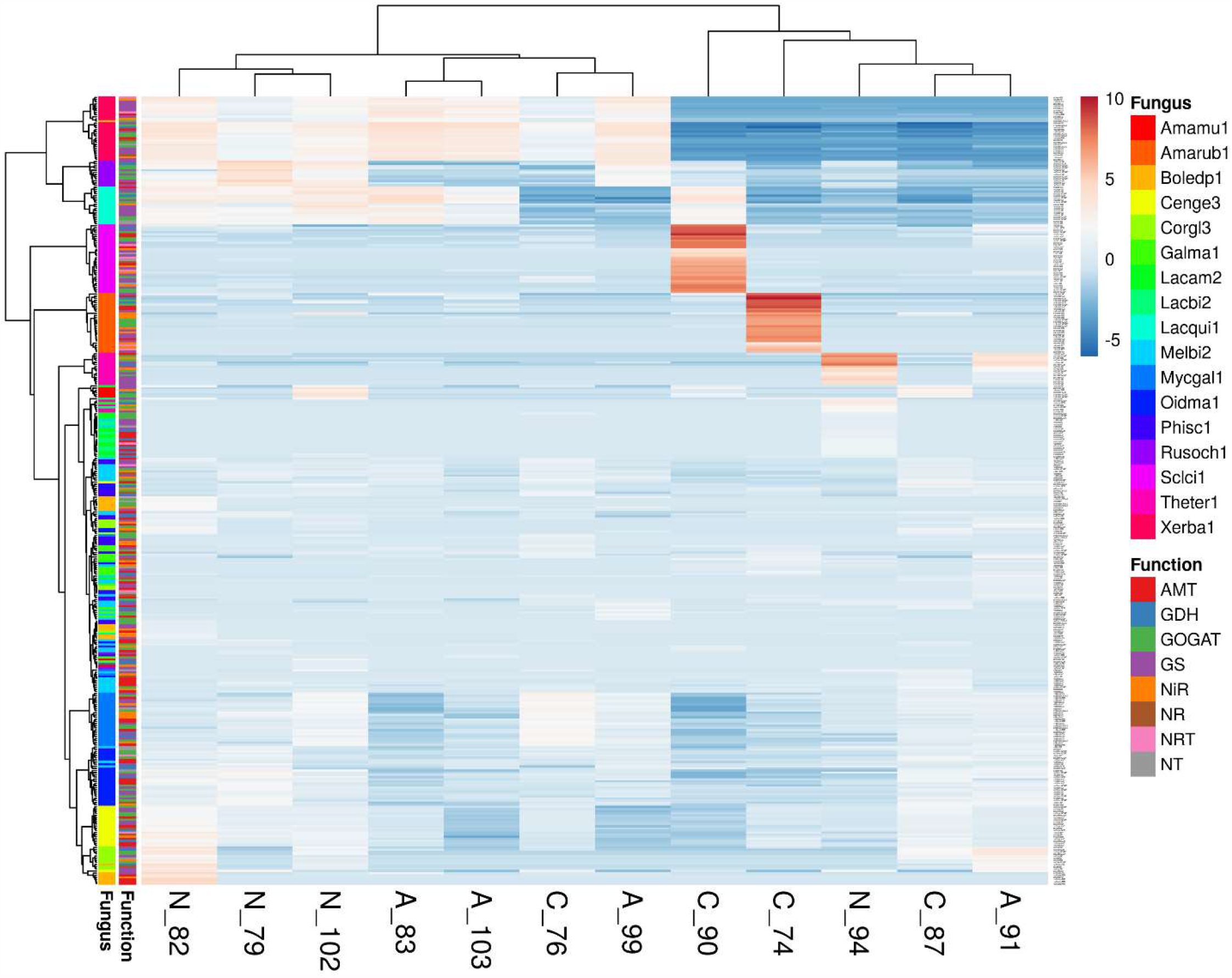
Cluster analysis of N-related transporters and enzymes represented by transcript abundances for ectomycorrhizal, endophytic and saprotrophic fungi colonizing beech roots. C = control, A = ammonium treatment, N = nitrate treatment, abbreviations for the fungi are shown in Table 1. Original values of the transcript levels were ln(x + 1)-transformed. Rows are centered; no scaling is applied to rows. Both rows and columns are clustered using Euclidean distance and Ward linkage. 369 rows, 12 columns.

### ^15^N application records strong N uptake by roots with increased root N concentrations

The EMRTs showed a strong ^15^N enrichment in response to ^15^NH_4_^+^ and ^15^NO_3_^-^treatment (Table 3) although specific effects related to mineral N provision were not discovered in the EMF metatranscriptome. The ^15^N enrichment in the root system decreased with increasing distance from the root tips and was about 2-times lower in fine roots, and about 6- to 8-times lower in coarse roots than in EMRTs (Table 3). The N content of the ^15^N-treated roots was slightly and significantly increased in comparison to control roots (Table 3) supporting that short-term N application caused enhanced N uptake. Thus, the N treatments triggered a significant decrease in the fine root C/N ratio compared to the controls (Table 3). The soil N content was not markedly affected by ^15^N application and the ^15^N signatures of nitrate- and ammonium-treated soils did not differ (Table 3). Overall, the beech root systems accumulated 1.5 ± 0.7% and 1.2 ± 0.6% of ^15^N from ammonium or from nitrate, respectively (Table 3). Since assimilation of inorganic nitrogen requires carbon skeletons (51), we measured fine root non-structural carbohydrate concentrations. However, no significant effects of N treatment on the carbohydrate concentrations were detected (Table 3).

**TABLE 3.** Biomass and root and soil chemistry in control and ^15^N-ammonium or ^15^N-nitrate treated cosms^c^

| Variable | Control | Ammonium | Nitrate | F value | P value |
| --- | --- | --- | --- | --- | --- |
| Biomass of CR (g cosm<sup>-1</sup>) | 3.44 ± 1.37 a | 2.87 ± 1.21 a | 3.40 ± 1.16 a | 0.5395 | 0.5905 |
| Biomass of FR (g cosm<sup>-1</sup>) | 0.88 ± 0.47 a | 0.66 ± 0.41 a | 0.70 ± 0.28 a | 0.6865 | 0.5138 |
| Biomass of EMF_RA (g cosm<sup>-1</sup>) | 0.32 ± 0.17 a | 0.23 ± 0.04 a | 0.20 ± 0.08 a | 0.0759 | 0.9275 |
| Soil dry mass (g cosm<sup>-1</sup>) | 1227 ± 202 a | 1149 ± 310 a | 1155 ± 466 a | 0.141 | 0.8693 |
| <sup>15</sup>N enrichment (mg g<sup>-1</sup> CR) | na | **0.11 ± 0.03 b** | **0.06 ± 0.02 a** | **9.8675** | **0.008512** |
| <sup>15</sup>N enrichment (mg g<sup>-1</sup> FR) | na | 0.27 ± 0.11 a | 0.21 ± 0.04 a | 1.1993 | 0.295 |
| <sup>15</sup>N enrichment (mg g<sup>-1</sup> EMRT) | na | 0.64 ± 0.45 a | 0.52 ± 0.11 a | 0.0285 | 0.8768 |
| <sup>15</sup>N enrichment (mg g<sup>-1</sup> soil) | na | 0.0171 ± 0.0062 a | 0.0237 ± 0.017 a | 0.8583 | 0.3725 |
| <sup>15</sup>N enrichment in roots (mg cosm<sup>-1</sup>) | na | 0.53 ± 0.25 a | 0.42 ± 0.22 a | 0.7395 | 0.4067 |
| <sup>15</sup>N enrichment in soil (mg cosm<sup>-1</sup>) | na | 18.35 ± 4.63 a | 20.76 ± 6.38 a | 0.6558 | 0.4338 |
| N (mg g<sup>-1</sup> CR) | 9.16 ± 2.28 a | 10.48 ± 2.29 a | 9.19 ± 1.77 a | 0.9419 | 0.4065 |
| N (mg g<sup>-1</sup> FR) | **12.90 ± 1.72 a** | **14.68 ± 1.60 ab** | **15.13 ± 1.36 b** | **4.5612** | **0.02334** |
| N (mg g<sup>-1</sup> EMRT) | 16.07 ± 4.83 a | 17.44 ± 0.06 a | 18.69 ± 1.85 a | 0.4571 | 0.6507 |
| N (mg g<sup>-1</sup> soil) | 4.34 ± 3.06 a | 3.98 ± 2.93 a | 4.61 ± 3.85 a | 4e-04 | 0.9996 |
| C (mg g<sup>-1</sup> CR) | **450.95 ± 5.74 ab** | **456.03 ± 8.04 b** | **444.97 ± 7.16 a** | **4.4774** | **0.02472** |
| C (mg g<sup>-1</sup> FR) | 479.61 ± 14.80 a | 472.13 ± 21.32 a | 467.36 ± 13.35 a | 1.1062 | 0.3502 |
| C (mg g<sup>-1</sup> EMRT) | 435.74 ± 93.60 a | 462.10 ± 1.65 a | 465.03 ± 9.07 a | 0.202 | 0.8217 |
| C (mg g<sup>-1</sup> soil) | 114.99 ± 88.14 a | 104.78 ± 82.78 a | 123.77 ± 113.81 a | 0.0699 | 9.9327 |
| C:N ratio in CR | 51.93 ± 12.41 a | 45.32 ± 10.40 a | 50.00 ± 9.54 a | 0.7282 | 0.4951 |
| C:N ratio in FR | **37.70 ± 4.75 b** | **32.38 ± 2.43 a** | **31.05 ± 2.20 a** | **7.8149** | **0.003106** |
| C:N ratio in EMRT | 28.12 ± 6.45 a | 26.54 ± 0.00 a | 25.02 ± 2.05 a | 0.3095 | 0.7434 |
| C:N ratio in soil | 25.75 ± 2.90 a | 25.78 ± 2.21 a | 25.42 ± 2.75 a | 0.0423 | 0.9586 |
| N-NH<sub>4</sub><sup>+</sup> (mg g<sup>-1</sup> FR) | 0.11 ± 0.03 a | 0.09 ± 0.03 a | 0.09 ± 0.03 a | 0.3329 | 0.7253 |
| N-NO<sub>3</sub><sup>-</sup> (mg g<sup>-1</sup> FR) | 1.96 ± 0.32 a | 2.59 ± 0.73 a | 1.87 ± 0.46 a | 2.2306 | 0.1634 |
| Glucose (mg g<sup>-1</sup> FR) | 16.99 ± 2.73 a | 16.32 ± 2.20 a | 16.36 ± 1.98 a | 0.1062 | 0.9003 |
| Fructose (mg g<sup>-1</sup> FR) | 9.06 ± 1.23 a | 8.51 ± 1.84 a | 7.71 ± 0.90 a | 0.9712 | 0.415 |
| Sucrose (mg g<sup>-1</sup> FR) | 0.61 ± 0.47 a | 0.51 ± 1.02 a | 0.82 ± 1.64 a | 0.0759 | 0.9275 |
| Starch (mg g<sup>-1</sup> FR) | 21.20 ± 11.95 a | 18.40 ± 8.66 a | 16.03 ± 5.20 a | 0.2411 | 0.7907 |
| TNSC (mg g<sup>-1</sup> FR) | 47.87 ± 14.25 a | 43.75 ± 12.51 a | 40.93 ± 6.33 a | 0.3484 | 0.7149 |

<sup>c</sup>Analyses were conducted two days after watering each cosm with 35 mg <sup>15</sup>N. Mean soil pH was 3.6 ± 0.1 and mean relative soil water content was 47.6 ± 28.9% (n = 25) across all studied cosms. Data in table shows means ± SD for dry samples. For dry mass: control (n = 9), ammonium (n = 8), nitrate (n = 8). For <sup>15</sup>N, C and N: control (n = 9), ammonium (n = 7), nitrate (n = 7), except for root tips where control (n = 5), ammonium (n = 2), nitrate (n = 3). For ammonium-N, nitrate-N and non- structural carbohydrates in FR, n = 4 per treatment. Significant differences among treatments (control, ammonium, nitrate) at p < 0.05 (Tukey HSD test) are shown in rows and marked in bold. Abbreviations: coarse roots (CR), fine roots (FR), ectomycorrhizal root tips (EMRTs), TNSC (total non-structural carbohydrates), na = not applicable because mean <sup>15</sup>N values of non-labeled controls were subtracted from the <sup>15</sup>N-treated samples.

### Beech transcriptome responds to nitrate and ammonium treatments activating N assimilation

Mapping of the RNA reads to the beech genome resulted in a total of 55,408 beech transcript ids or gene models before normalization (Data set 4) and 27,135 beech gene models after normalization (Data set 5) in DESEq2. Ammonium and nitrate treatment resulted in 75 and 74 differentially expressed beech gene models (DEGs), respectively, with both treatments sharing 26 DEGs (Fig. 4A) and indicating overlapping responses to ammonium and nitrate. Among these overlapping DEGs, a putative glutamine synthetase (*GS*, AT5G35630.2) showed the highest upregulation, along with five putative cysteine-rich receptor-like protein kinase orthologs of *Arabidopsis thaliana* (*CRK8*, AT4G23160.1), outward rectifying potassium channel protein (*ATKCO1*, AT5G55630.2), HXXXD-type acyl transferase family protein (AT5G67150.1), hemoglobin 1 (*HB1*, AT2G16060.1), molybdate transporter 1 (*MOT1*, AT2G25680.1), and early nodulin-like protein 20 (*ENODL20*, AT2G27035.1) (Fig. 4B). Moreover, among the downregulated overlapping DEGs were a cinnamate-4-hydroxylase (*C4H*, AT2G30490.1) which plays a role in plant phenylpropanoid metabolism, growth, and development (61), eight orthologs coding for a DNAse 1-like superfamily protein (AT1G43760.1), AP2/B3-like transcription factor family proteins (*VRN1*, AT3G18990.1) which are involved in regulation of the vernalization pathway (62, 63), subtilase family protein (AT5G45650.1), Ankyrin repeat family protein (AT3G54070.1), LRR and NB-ARC domains-containing disease resistance protein (*LRRAC1*, AT3G14460.1) known to play roles in the immune response against biotrophic fungi and hemibiotrophic bacteria (64), and NB-ARC domain-containing disease resistance protein (AT4G27190.1) (Fig. 4B).

**FIG 4.**
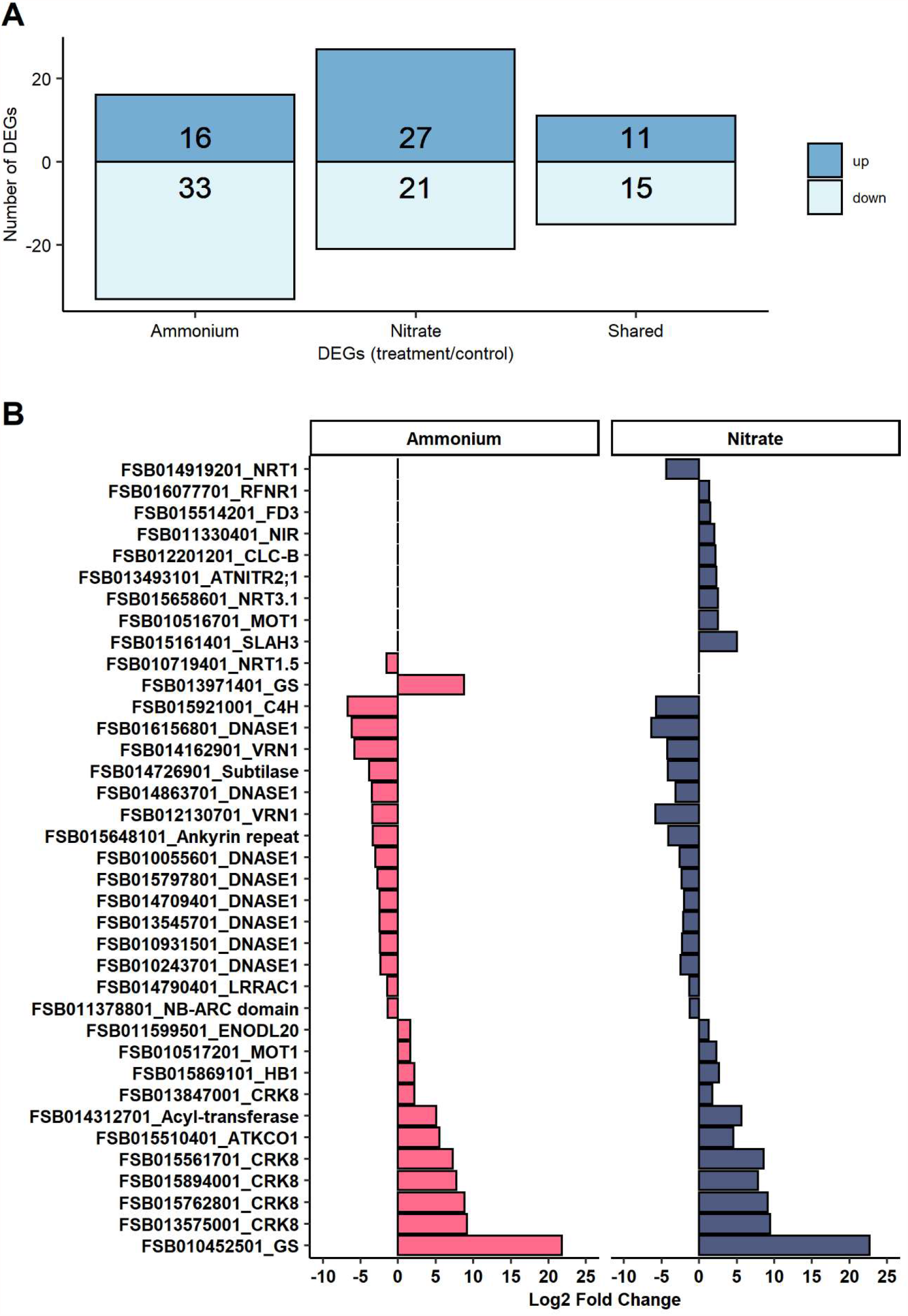
Differentially expressed genes (DEGs) in response to ammonium or nitrate exposure. A) Number of unique and shared DEGs (padj < 0.05 and 2-fold change) in response to ammonium or nitrate treatment. B) Log2-fold changes of shared DEGs and DEGs related to N-metabolisms in beech roots in response to increased ammonium or nitrate treatment relative to control conditions (n = 4 per treatment). The complete information, gene model ID and names are provided in Data set 5.

Among unique responses to ammonium treatment were upregulation of a further *GS* ortholog (AT5G35630.2) and downregulation of a putative nitrate transporter gene (*NRT1*.*5*, AT1G32450.1) (Fig. 4B) known to load nitrate into the xylem and to be induced at high or low nitrate concentration in *A. thaliana* (65). Among the unique DEGs detected in response to nitrate treatment and known to play roles in nitrate translocation and metabolism were a putative high affinity nitrate transporter (*NRT3*.*1*, AT5G50200.1) which was upregulated along with a putative nitrite transmembrane transporter (*ATNITR2;1*, AT5G62720.1, see (66), a nitrite reductase 1 (*NIR*, AT2G15620.1), a molybdate transporter 1 (*MOT1*, AT2G25680.1), a SLAC1 homologue 3 (*SLAH3*, AT5G24030.1), and chloride channel b (*CLC-B*, AT3G27170.1) genes (Fig. 4B). Furthermore, transcripts for the root-type ferredoxin:NADP(H) oxidoreductase gene (*RFNR1*, AT4G05390.1) which supplies electrons to Ferredoxin-dependent enzymes (e.g., Fd-NiR and Fd-GOGAT) (67), and a ferredoxin 3 gene (*FD3*, AT2G27510.1) which enables electron transfer activity were also upregulated, while a putative nitrate transporter gene (*NRT1/ PTR FAMILY 6*.*2*, AT2G26690.1) was down regulated (Fig. 4B). Other genes involved in N assimilation exhibited basal transcript levels, including those coding for the enzymes GOGAT and GDH detected under nitrate, ammonium, and control conditions but not differentially regulated.

Classification of beech DEGs into Mapman bins revealed a significant overrepresentation of genes involved in “nitrogen metabolism” for both ammonium and nitrate treatments (Fig. 5). Significantly overrepresented metabolic processes for the nitrate treatment included “oxidative pentose phosphate pathway (OPP),” “protein,” “redox,” “secondary metabolism,” “signaling” and “stress” (Fig. 5). For the ammonium treatment, significantly overrepresented functions included “DNA,” “hormone metabolism,” “secondary metabolism,” “signaling,” “stress” and “transport” (Fig. 5). Pathway enrichment analysis of Gene Ontology terms of beech DEGs in g:Profiler returned significant results for nitrate, but not for ammonium treatment. DEGs from the nitrate treatment resulted in 38 significantly enriched GO terms involving nitrate-related molecular level functions and four biological processes including “nitrate transmembrane transporter activity,” “nitrite reductase activity,” “response to nitrate” and “nitrate transport” (Table S3). Plant immune responses induced by nitrate were also evident via the enrichment of a putative isochorismate synthase gene (*ICS2*, AT1G18870) and a flavin-dependent monooxygenase 1 gene (*FMO1*, AT1G19250). ICS2 is involved in the biosynthesis of vitamin K_1_ (68) and potentially in salicylic acid biosynthesis (69, 70). *FMO1* is involved in the catalytic conversion of pipecolic acid to N-hydroxypipecolic acid (NHP) which plays a role in plant acquired systemic resistance to infection by pathogens (71).

**FIG 5.**
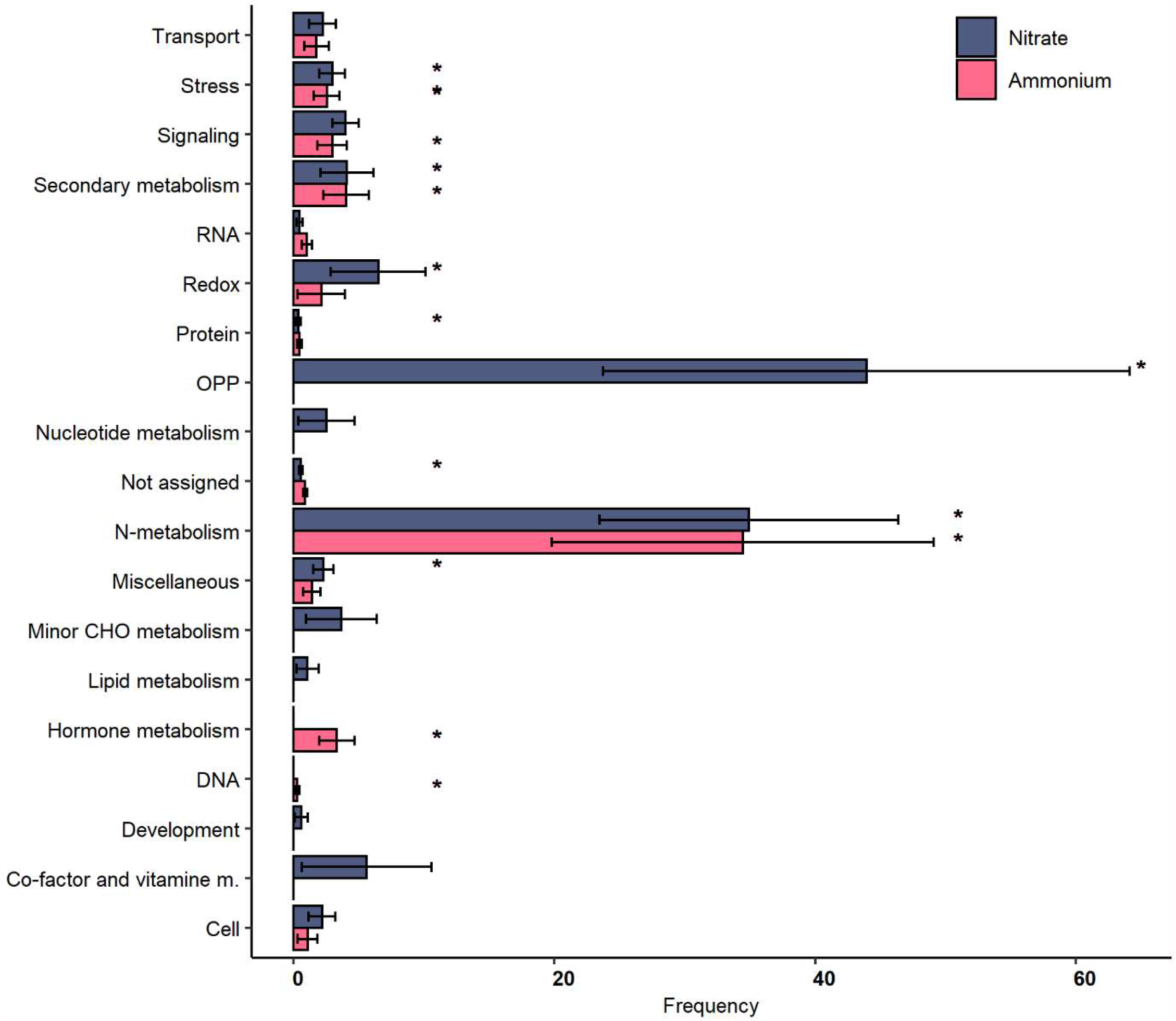
Classification of beech root DEGs in response to nitrate or ammonium treatment. Genes were classified according to Mapman bins using Classification superviewer in BAR (http://bar.utoronto.ca/ntools/cgi-bin/ntools_classification_superviewer.cgi). Bins that were statistically significantly enriched are marked by an asterisk.

## DISCUSSION

### Ectomycorrhizal and root acquisition of nitrate or ammonium

A central aim was to gain insights into gene regulation in self-assembled ectomycorrhizas by targeting the transcriptomes of the fungi living in active symbiosis with beech roots in response to N provision. To challenge fungal metabolism, we applied N treatments that caused about 3- and 14-fold increases in the available NH_4_^+^-N (about 15.1 µg g^-1^ soil DW) and NO_3_^-^-N (about 2.3 µg g^-1^ soil DW), respectively. The magnitude of these variations was similar to temporal fluctuations of NH_4_^+^-N and NO_3_^-^-N observed in soil of beech stands with 2-fold and 10-fold changes for NH_4_^+^-N and NO_3_-N, respectively (72). Therefore, it was expected to elicit representative N responses in the naturally assembled EMF communities. The EMF assemblages in our study showed the typical patterns known for temperate beech forests with high diversity (21, 23, 24), dominance of certain species (73, 74) (e.g., genus *Amanita, Xerocomus* and *Scleroderma* in this study), and non-uniform occurrence in the tree roots. However, the two-day N treatments were not expected to affect the fungal community structure because colonization and establishment of new ectomycorrhizas takes weeks or months rather than days (75, 76), and shifts in fungal communities towards more nitrophilic fungi occur as a consequence of long-term exposure to high N loads (77–81).

Our EMF community were composed of taxa characteristic for acidic sandy, nutrient poor soils, including genera in the orders Agaricales, Boletales, Russulales, Helotiales, Myltinidales and Thelephorales (Data set 1). These fungi vary in their foraging strategies being equipped with different types of hyphae for scavenging N. *C. geophilum* which is the most widespread fungus and known for its tolerance to drought (82) produces short and medium-distance hyphae, while the hyphae of *Amanita* (medium-distance smooth or long-distance), *Cortinarius* (medium-distance fringe), *Laccaria* and *Telephora* (medium-distance smooth), *Lactarius* (contact, short and medium distance), *Russula* (contact), and *Scleroderma* and *Xerocomus* (long-distance) are also diverse (78, 83). EMF that produce hydrophilic hyphae of contact, short, and medium-distance smooth exploration types were reported to respond positively or to display a mixed response to mineral N enrichment, whereas EMF with medium-distance fringe hydrophobic hyphae are the most sensitive, and those with long-distance hydrophobic hyphae vary in their responses to mineral N (78).

The availability of nitrate and ammonium ions in the soil of this experiment was made highly dynamic by a sudden increase. The high mobility of the negatively charged nitrate ions in the soil solution make it more available for the roots than the positively charged ammonium ions which tend to be fixed by soil colloids, and while both ions are prone to leaching in sandy soils, ammonium retention by organic matter and clay minerals is generally higher (5, 13, 14, 84). The lower energy cost needed for ammonium metabolism make its utilization more advantageous than nitrate. This was previously observed in EMF (30–35), and in agreement, we found higher translocation of ^15^N from NH_4_^+^ than from NO_3_^-^to the coarse roots. Enrichment of the newly applied ^15^N in the EMRTs was strong but did not differ between N form applied. We cannot exclude ammonification by soil microbes potentially converting NO_3_^-^ to NH_4_^+^ in the soil before its uptake by the EMF, thus contributing to similar ^15^N accumulation patterns in the ectomycorrhizas after NO_3_^-^ or NH_4_^+^ application. Microbial turnover rates are estimated to be about 24 h for ammonium and a few days for nitrate (85). However, the significant transcriptional regulation of nitrate marker genes in beech roots under nitrate exposure supports that NO_3_^-^ was taken up by the root system. In fine root cells, NO_3_^-^was considerably more abundant than NH_4_^+^ as observed in beech trees in field conditions (72, 86) and unaffected by N addition. Our results demonstrate that the newly acquired ^15^N was metabolized because the root N concentrations increased but not the levels of ammonium or nitrate.

### N assimilation uncovers fungal taxon-specific but not N-induced transcription patterns in root associated fungal communities

Despite the compelling support for N uptake and assimilation in roots, the EMF metatranscriptome did not show any significant changes related to N metabolism. Initially, we hypothesized that if the root-associated fungi and the beech root cells responded like a synchronized “superorganism,” both fungi and roots would show similar patterns in transcriptional regulation. However, this hypothesis is rejected because N-responsive DEGs were found in beech but not in the EMF metatranscriptomes, except for KOG4381 and KOG4431 which were induced by nitrate. Closer inspection revealed that KOG4381 occurred only in *Thelephora terrestris* and *Russula ochroleuca*, thus not reflecting a community response and rather suggesting that in the symbiotic system, the host and EMF partners respond as individual autonomous units. KOG4431 was present in nine EMF, including *Laccaria amethystina, Meliniomyces bicolor, Russula ochroleuca, Scleroderma citrinum, Thelephora terrestris, Xerocomus badius, Amanita rubescens, Boletus edulis* and *Cenococcum geophilum*. Further analyses are needed to clarify the role of these two KOGs in nitrate signaling. Although differentially expressed KOGs were limited in the fungal metatranscriptomes, putative transporters (NRT/NIT, AMT) and enzymes (NR, NiR, GS, GOGAT, and GDH) were transcribed (Fig.6), representing all necessary steps for mineral N uptake and assimilation into amino acids. In controlled laboratory studies, many of these transporters and enzymes have been characterized in EMF and were regulated by N form and availability. For instance, high affinity nitrate/nitrite transporters (NRT2), nitrate reductase (NR) and nitrite reductase (NiR1) in *Hebeloma cylindrosporum* (87, 88), NRT2, NR1 and NiR1 in *Tuber borchii* (89, 90), NRT, NR and NiR in *L. bicolor* (44, 91), high and low affinity ammonium transporters (AMT1, AMT2, and AMT3) in *H. cylindrosporum* (50, 92), AMT2 in *Amanita muscaria* (93), and AMT1, AMT2 and AMT3 in *L. bicolor* (44). Although we did not find N-induced regulation of specific genes, GO term enrichment analysis shows that functions related to N assimilation and carbon metabolism were enriched across all studied EMF. We suggest that at the whole EMF community level, the primary metabolism is genetically equipped for handling fluctuating environmental N availability and host-derived C supply.

The observed stability of the fungal metatranscriptome was unexpected because stable isotope labeling and electrophysiological studies showed distinct responsiveness of different fungal taxa to environmental changes in self-assembled communities (27, 28, 94) and controlled studies (cited above and in the introduction) showed significant regulation of N-related genes. Our study does not exclude that there were N-induced responses in distinct fungi, but weak effects might have been masked by the variability of EMF species occurrence in individual cosms. Presumed species-specific responses to N fertilization were probably also overridden by inter-specific differences. This can be inferred from the observation that arrays of N-related genes clustered quite strictly according to species but not according to the genes with similar functions. Our identification of expression patterns for the fungi under study is an important, novel result underpinning trait stability within naturally-assembled EMF in beech roots.

### Ammonium and nitrate induce specific assimilation patterns in beech roots

Our initial hypothesis was that EMF shield the plant cells against major fluctuations in N availabilities and therefore we expected no or moderate changes in the beech root transcriptome after N fertilization. This hypothesis is rejected since both ammonium and nitrate treatments caused drastic changes in the beech root transcriptome. The strategy of European beech for dealing with high loads of inorganic N availability was transcriptional upregulation of genes involved in N uptake and assimilation, as observed in *Arabidopsis* (51),a non-mycorrhizal species. The transcription patterns in response to nitrate and ammonium were clearly distinguishable, in agreement with other studies that documented nitrate- and ammonium-specific effects on gene regulation, signaling and lateral root growth (95–100). Notably, transcripts belonging to putative NRTs and to enzymes (NR, NiR, GS) were significantly upregulated in the nitrate treatment encompassing the suite of reactions required for NO_3_^-^reduction and incorporation into amino acids (Fig. 6). In addition, upregulation of root ferredoxin and molybdate transporters pointed to an enhanced need for reducing power and the biosynthesis of NR, which requires molybdate in its active center (101). The significant activation of defenses against biotrophic fungi by nitrate was also remarkable. Similar results were shown in leaves of non-mycorrhizal nitrate-fed trees (102). In the ammonium treatment, significant upregulation of transcripts levels for two GS enzymes was detected, while a NRT1.5, potentially loading nitrate into the xylem, was downregulated. Remarkably, nitrate and ammonium treatments showed a common pattern with strong upregulation of *GS* and *CRK*-like genes (Fig. 4B). CRK receptor kinases are involved in stress, plant pathogen response and cell death (103, 104). The *Arabidopsis* ortholog CRK8 is regulated in senescing leaves (105), and while a function in N metabolism appears likely, controlled experiments are needed. Overall, these results were in line with the expectation that nitrate-specific, ammonium-specific and overlapping responses were to be found. We demonstrate for the first time that excessive N in EMRTs is actively metabolized by the plant. It remains unknown if NO_3_^-^ and NH_4_^+^ were taken up by EMF and transferred to the plant for further assimilation or if excessive N circumvented the fungal barrier, entering the plant directly (Fig. 6).

**FIG 6.**
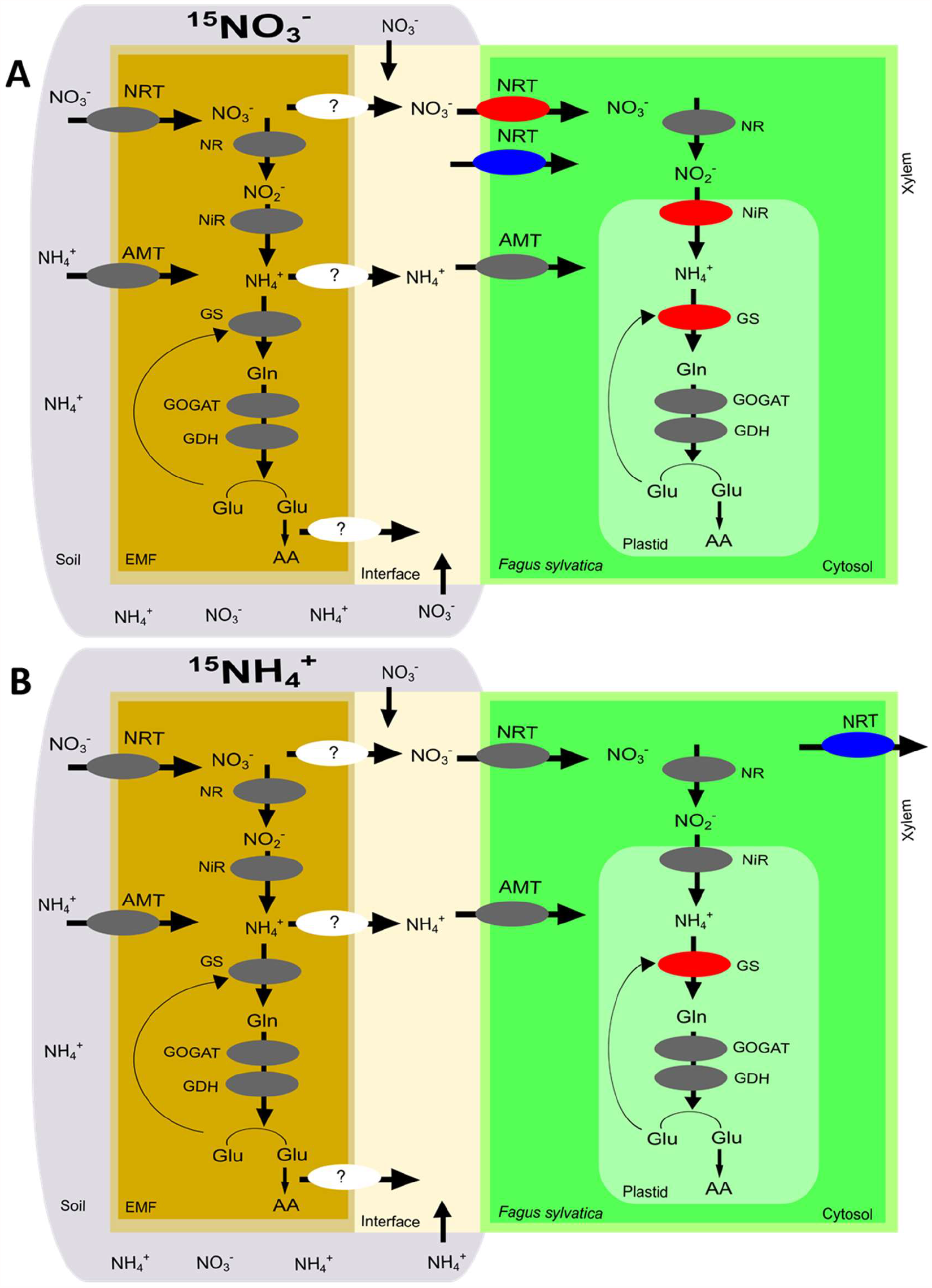
Scheme of the pathway for N uptake and assimilation in EMF and *Fagus sylvatica* based on transcription profiles. Regulation of ectomycorrhizal fungi and host transcripts encoding for transporters and enzymes involved in N uptake and assimilation detected in the nitrate treatment (A) and in the ammonium treatment (B). NRT: nitrate/nitrite transporter, NR: nitrate reductase, NiR: nitrite reductase; AMT: ammonium transporter, GS: glutamine synthetase, GOGAT: glutamate synthase; GDH: glutamate dehydrogenase. Gray: detected but not regulated, Red: upregulated, Blue: downregulated, White: not detected.

In conclusion, effects of high levels of ammonium or nitrate were not evident in the EMF metatranscriptome, whereas the host tree responded to ammonium and to nitrate by upregulating genes involved in assimilation of the surplus inorganic N into organic forms. Although it is unknown whether the applied ^15^N sources underwent conversions due to microbial activities, the response of European beech indicated that a significant proportion of ammonium and nitrate was taken up in the originally added form. The fungal transcriptomes suggested species-specific N-metabolic responses, implying significant trait stability for N turnover and suggesting that EMF in temperate beech forests are resistant to short-term fluctuations in environmental N. However, further work is required to investigate to what extent this tolerant capacity can be sustained and its ecological relevance under chronic N exposure.

## MATERIALS AND METHODS

### Tree collection, maintenance, and experimental setup

European beech (*Fagus sylvatica* L.) saplings were collected on March 7^th^, 2018, in a 122-year-old beech forest (53°07’27.7”N, 10°50’55.7”E, 101 m above sea level, Göhrde, Lower Saxony, Germany). The soil type is podzolic brown earth with parent material consisting of fluvio-glacial sands (106). In 2017, the mean annual temperature was 9.9 °C and the total annual precipitation 768 mm, whereas on the day of tree collection the mean air temperature was 4.6 °C and the precipitation was 0.66 mm (https://www.dwd.de). The beech saplings (n = 34) were excavated using polyvinyl chloride cylinders (diameter: 0.125 m, depth: 0.2 m), which were placed around a young tree, hammered into the ground to a depth of 0.2 m, and then carefully lifted to keep the root system in the intact forest soil. These experimental systems are referred to as cosms. The cosms were transported to the Forest Botanical Garden, University of Goettingen (51°33’27.1”N 9°57’30.2”E) where they were maintained outdoors under a transparent roof and exposed to natural climatic conditions except for rain (Table S4). A green shading net was placed over the roof to protect the trees from direct sun similar as in the forest. Thereby, on average, the full sun light was reduced on sunny days from 1125 µmol m^2^ s^-1^ to 611 µmol m^-2^ s^-1^ PAR (photosynthetically active radiation) and on cloudy days from 284 µmol m^-2^ s^-1^ to 154 µmol m^-2^ s^-1^ (Quantum/Radiometer/Photometer model 185B, LI-COR Inc., Lincoln, NE, USA). The cosms were regularly watered with demineralized water. Control of the water quality (flow analyzer, SEAL AutoAnalyzer 3 HR, SEAL Analytical GmbH, Norderstedt, Germany) revealed 0.2 mg NH_4_^+^ L^-1^ and no detectable NO_3_^-^ in the irrigation water. The cosms were randomly relocated every other day to avoid confounding positional effects. The trees were grown under these conditions until July 2018. By this time, the trees had a mean height of 0.401 ± 0.08 m and a root collar diameter of 6.11 ± 0.95 mm. The trees were about 8 (± 2) years old based on the number of growth scars along the stem (107). Before the ^15^N treatments, ammonium and nitrate were measured in the soil (details in Text S1). The cosms contained 15.1 ± 11.3 µg NH_4_^+^-N g^-1^ soil DW and 2.3 ± 1.3 µg NO_3_^-^-N g^-1^ soil DW (n = 3, ±SD), equivalent to approximately 9.1 mg NH_4_^+^-N and 1.8 mg NO_3_^-^-N cosm^-1^.

### Application of ^15^N-labelled ammonium and nitrate

Before labeling, even distribution of irrigation solution in the soil was tested on separate cosms using blue dye (“GEKO” Lebensmittelfarbe, Wolfram Medenbach, Gotha, Germany) in water. The experimental cosms were assigned the following treatments: control (no nitrogen application), ^15^NH_4_^+^ application or ^15^NO3^-^application. The cosms were surface-irrigated at 7 am with 60 ml of either 19.85 mM ^15^NH_4_Cl (99% ^15^N, Cambridge Isotope Laboratories, Inc., MA, US, pH 5.47) or a 19.98 mM ^15^KNO_3_ (99% ^15^N, Cambridge Isotope Laboratories, pH 6.23) solution prepared in autoclaved deionized water. Controls were irrigated with 60 ml autoclaved demineralized water (pH 6.07). Each of these treatments were repeated the next day, resulting in a total application of 35.96 mg ^15^N in the nitrate treated cosms or 35.74 mg ^15^N in the ammonium treated cosms, corresponding to mean additions of approximately 30 µg ^15^N g^-1^ dry soil. Treatments were conducted in two batches: batch 1: ^15^N application on July 17^th^, 2018 and harvest on July 19^th^, 2018, n = 9 cosms; batch 2: ^15^N application on July 31^st^, 2018 and harvest on August 2^nd^, 2018, n = 16 cosms.

### Cosm harvest

The cosms were harvested 48 h after initial ^15^N application. The tree-soil compartment was pushed out of the cylinder, collecting all parts. Roots were briefly rinsed with tap water, then with deionized water, and gently surface-dried with paper towels. The root tips were clipped off, shock-frozen in liquid nitrogen, and stored at -80°C. Aliquots of fine roots were shock-frozen in liquid nitrogen and stored at -80°C and -20°C and soil aliquots at -20°C. During the harvests, the fresh masses of all fractions (leaves, stem, coarse roots, fine roots, root tips and soil) were recorded and aliquots were taken for dry-to-fresh mass determination after drying at 40°C (leaves, stems, soil) or after freeze-drying (coarse roots, fine roots and root tips). Biomass and soil mass in the cosms were calculated:

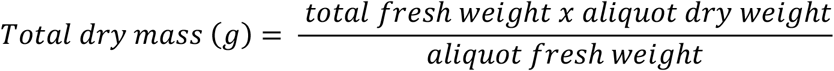

### Soil and root chemistry

Soil pH was measured with a WTW pH meter 538 (WTW, Weilheim, Germany) using a ratio of dry sieved soil to water of 1: 2.5 according to the Forestry Analytics manual (108), A3.1.1.1, page 2. The water content in the soil was calculated as:

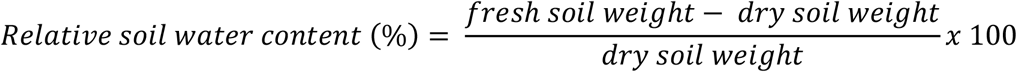

For ^15^N analyses, freeze-dried aliquots of soil, root tips, fine and coarse roots were milled using a ball mill (Type MM400, Retsch GmbH, Haan, Germany) in stainless steel grinding jars at a frequency of 30/sec in 20 second intervals to avoid heating the sample. The powder (control samples: 1.5 to 2 mg plant tissues, 5 mg soil; labeled samples: 1.5 to 3 mg plant tissue, 5 to 13 mg soil) was weighed into tin capsules (IVA Analysentechnik GmbH & Co KG, Meerbusch, Germany) and measured at the KOSI (Kompetenzzentrum Stabile Isotope, Göttingen, Germany). The ^15^N samples were measured in the isotope mass spectrometer (Delta V Advantage, Thermo Electron, Bremen Germany) and an elemental analyzer (Flash 2000, Thermo Fisher Scientific, Cambridge, UK) and the non-labeled control samples in a mass spectrometer: Delta plus, Finnigan MAT, Bremen, Germany and elemental analyzer: NA1110, CE-Instruments, Rodano, Milano, Italy). Acetanilide (10.36 % N, 71.09 % C, Merck KGaA, Darmstadt, Germany) was used as the standard. Enrichments of ^15^N in the ectomycorrhizal root tips (EMRTs), fine roots, coarse roots and soil were calculated as:

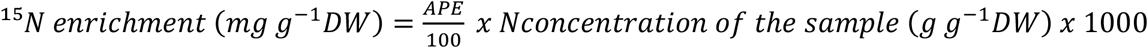

where

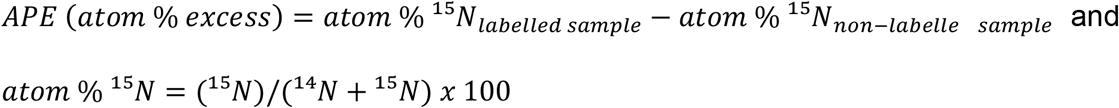

For determination of NH_4_^+^, NO_3_^-^ and non-structural carbohydrates, frozen fine roots (−80°C) were milled (MM400, Retsch GmbH) under liquid nitrogen to avoid thawing. For mineral N determination, the frozen powder (approximately 55 mg per test) were extracted as described before (109) with slight modifications and measured spectrophotometrically with the Spectroquant® (1.09713.0002) Nitrate and Ammonium (1.14752.0002) Test Kits (Merk, KGaA, Darmstadt, Germany). Glucose, fructose, sucrose and starch were extracted from approximately 75 mg root powder and were measured enzymatically as described before (110). Details of all procedures are reported in (Text S1).

### DNA extraction, Illumina sequencing, bioinformatic processing and data analyses of fungi

Root tips (from -80°C) were homogenized in liquid nitrogen using sterilized mortar and pestle. Each powdered, frozen sample was split in two parts: one for DNA extraction and Illumina sequencing of the fungal ITS2 gene and the other for RNA extraction and mRNA sequencing. DNA was extracted from approximately 200 mg powder of root tips using the innuPREP Plant DNA Kit (Analytik Jena, AG, Jena, Germany). Extraction, purification, processing and sequencing are described in detail (Text S1). Briefly, the fungal nuclear ribosomal DNA internal transcribed spacer (ITS2) region was amplified by Polymerase Chain Reaction (PCR) using the primer pair ITS3_KYO2 (111) and ITS4 (112), both containing specific Illumina overhang adapters (in italics, primers underlined): forward (Miseq_ITS3_KYO2): 5’-*TCGTCGGCAGCGTCAGATGTGTATAAGAGACAG*<u>GATGAAGAACGYAGYRAA</u>-3’; reverse (Miseq_ITS4): 5’-*GTCTCGTGGGCTCGGAGATGTGTATAAGAGACAG*<u>TCCTCCGCTTATTGATATGC</u>-3’

After the PCR, the amplicons were purified and sequenced on a MiSeq flow cell using Reagent Kit v3 and 2×300 pair-end reads (Illumina Inc., San Diego, USA) according to the manufacturer’s instructions at the Göttingen Genomics Laboratory (G2L, Institute of Microbiology and Genetics, Georg-August-University Göttingen, Göttingen, Germany). The raw sequences were quality filtered and clustered according to amplicon sequence variants (ASVs) at 97% sequence identity (i.e., operation taxonomic units, OTUs). Fungal reads were mapped to operational taxonomic units (OTUs) and abundance tables were generated. Taxonomic assignment of OTUs was carried out against the UNITE database v8.2 (04.02.2020),(113). All unidentified ASVs were searched (blastn) (114) against the nt database (2020-01-17) to remove non-fungal ASVs. Extrinsic domain ASVs and unclassified ASVs were discarded from the taxonomic table. The fungal OTUs were assigned to trophic modes using the FUNGuild annotation tool (115). The sequencing depth per sample was controlled by rarefaction analysis using the package ampvis2 (116) and the samples were normalized by rarefying to the sample with the lowest sequencing depth (i.e. 20,051 sequence reads). An overview of the sequence processing results is provided in Table S5, and the rarefied abundance table with taxonomic and guild assignment of OTUs is provided in (Data set 1).

### RNA extraction, library preparation, sequencing, and bioinformatic processing of the fungal metatranscriptome and beech transcriptome

Total RNA was isolated from the frozen powder of beech root tips using a CTAB method (117). The details have been reported in (Text S1). RNA integrity numbers (RIN), library preparation and sequencing were conducted at Chronix Biomedical GmbH (Goettingen, Germany). Twelve samples with RIN ranging from 6.7 to 7.9 were selected for polyA mRNA library preparation (Table S6). Libraries were constructed with the NEBNext RNA Ultra II Library Prep Kit for Illumina (New England Biolabs, Ipswich, Massachusetts, United States) from 1 µg of purified RNA according to the manufacturer’s instructions. Single-end reads with a length of 75 bp were sequenced on a NextSeq 500 Sequencing System instrument (Illumina, San Diego, CA, USA) with a sequencing depth of 100 million reads per sample.

Processing (trimming, quality filtering and adapter removal) of the raw sequence data (ca. 110 million reads per sample) resulted in approximately 109 million reads per sample (Table S6). The reads were mapped against the reference transcriptomes of *Fagus sylvatica* and 17 fungal species belonging to the same genera as those detected by ITS barcoding (Table 1). Reference beech sequences and annotations were downloaded from beechgenome.net (57) and reference fungal sequences and annotations were downloaded from the JGI MycoCosm database (56). The resulting 18 fasta files were concatenated to one single file, which was used to create an index file with bowtie2-build (118). The reads were mapped against this index file using bowtie2, resulting in one count table containing the reads for beech and fungi. On average, 61 % of the reads could be mapped (45 % to beech and 16 % to fungi) (Table S6). The raw count table was split into a beech transcriptome count table and fungal transcriptome count table. Normalization of the raw count tables and differential expression analyses relative to the control was conducted using the DESeq2 package (119), implemented in R (120). Differential expression analysis of the fungi was performed at the metatranscriptome level (i.e., the fungal raw count tables were aggregated by their EuKaryotic Orthologous Groups of protein identifiers (KOGs, https://img.jgi.doe.gov/)), dropping taxon-specific information for the gene models. Two fungal metatranscriptomes were considered: the full list fungi metatranscriptome (17 fungi) or the ectomycorrhizal-specific metatranscriptome (13 fungi). Gene models (European beech) or KOGs (fungal metatranscriptomes) with a Benjamini-Hochberg adjusted false discovery rate P_adjusted_ < 0.05 (121) and at least 2-fold change were considered as significant differentially expressed gene models (DEGs) or significant KOGs. The Enzyme Commission numbers assigned to the ectomycorrhizal fungal metatranscriptome were mapped to the Kyoto Encyclopedia of Genes and Genomes (KEGG) metabolic pathways against *L. bicolor* in KEGG Mapper (122). Functional enrichment analysis of fungal expressed genes was carried out in g:Profiler (123) against KEGG metabolic pathways with *Aspergillus oryzae* as reference since the model ectomycorrhizal fungus *L. bicolor* was not available. Since a main interest in our experiment was to obtain information on fungal N uptake and metabolism, we manually searched the complete fungal transcriptional database (Data set 2) for N-related transporters and enzymes using the key words “nitrate transporter,” “nitrate reductase,” “nitrite transporter,” “nitrite reductase,” “ammonium transporter”, “glutamine synthetase,” “glutamate synthase” and “glutamate dehydrogenase.” These terms were searched in the definition lines accompanying the annotations of each of the fungal transcripts: “kogdefline” = definition of the KOG identifiers, “ECnumDef” = definition of EC number, “iprDesc” = description of the InterPro identifiers, and “goName” = description of the Gene Ontology term. Cluster analyses was done in Clustvis (124). Gene Ontology term enrichment analysis of beech DEGs was also performed in g:Profiler (123). In addition, over-representation analysis of biological pathways based on the MapMan bin classification (Ath_AGI_LOCUS_TAIR10_Aug2012) of beech DEGs was performed using the Classification SuperViewer Tool (125) from the Bio-Analytic Resource for Plant Biology (http://bar.utoronto.ca/).

### Statistical analyses

The fungal community data were Hellinger-transformed and fitted into a non-metric multidimensional scaling (nMDS) ordination based on Bray-Curtis dissimilarity using the ‘vegan’ package version 2.5-6 (126) and ‘ggplot2’ function (127) in the R software (128). Permutational analysis of variance (‘adonis 2’) was used to test if the treatments resulted in significant effects on the fungal community or transcript composition. Quasi-Poisson regression models were used for over-dispersed count data (e.g., species richness) and general linear models were applied to normal distributed data, followed by Tukey’s HSD post-hoc test with the ‘multcomp’ package (129). When necessary, the data was transformed to meet normal distribution. If not indicated otherwise, data are shown as means (± SD). Linear regression analysis was conducted in R (128). One cosm from ammonium and one from nitrate treatment were excluded from the ^15^N analyses since the measured ^15^N values in soil were higher than the amount of added ^15^N.

## Supporting information

Rivera P&eacute;rez et al 2021 supplementary material

## Data availability statement

Raw sequences from the fungal ITS2 gene metabarcoding-Illumina sequencing are available in the Sequence Read Archive from the National Center for Biotechnology Information under BioProject accession number PRJNA736215 (130). Raw read data from RNA seq are also available at the ArrayExpress database under accession number E-MTAB-8931 (131). Additional supporting data (Data set 1-6) are accessible in Dryad (132).

## SUPPLEMENTAL MATERIAL

Table S1, Table S2, Table S3, Table S4, Table S5, Table S6, Figure S1, Text S1

## SUPPORTING DATA

Data_set_1_rarefied_fungal_otu_table.xlsx

Data_set_2_raw_rna_counts_fungi.xlsx

Data_set_3_normalized_rna_counts_fungal_metatranscriptomes.xlsx

Data_set_4_raw_rna_counts_fagus.xlsx

Data_set_5_normalized_rna_counts_fagus.xlsx

Data_set_6_plant_nutrients_and_environmental_conditions.xlsx

## ACKNOWLEDGEMENTS

This research was funded by the German Research Foundation (DFG) through the Research Training Group 2300 “Enrichment of European Beech Forests with Conifers” (contract number: 316045089, project SP4). We thank the Göhrde State Forest management office for authorizing tree collection in the forest. CARP is grateful to Dr. Serena Müller for assistance coordinating field work, to Michael Reichel, Ronny Thoms and Jonas Glatthorn for help collecting the trees in the forest, and Gaby Lehmann for help measuring ammonium/nitrate in soil.

CARP: Conceptualization, Methodology, Project administration, Investigation, Formal analysis, Writing – original draft, Writing – review & editing. DJ, DS, RD: Data curation, Writing – review & editing. AP: Conceptualization, Methodology, Formal analysis, Supervision, Writing – review & editing.

We declare no competing interests.

## Notes

### Competing Interest Statement

The authors have declared no competing interest.

