## Supplementary material for "Transcriptional Landscape of Ectomycorrhizal Fungi and Their Host Provide Insight into N Uptake from Forest Soil": Rivera P&eacute;rez et al 2021 supplementary material

**TABLE S1** OTU-level richness and diversity indices from the beech root-associated fungal community[a]

| Sample | Treatment | Taxa S | Dominance D | Simpson 1-D | Shannon H | Evenness e^H/S | Equitability J | Fisher alpha | Berger-Parker | Chao-1 |
|---|---|---|---|---|---|---|---|---|---|---|
| C_74 | demineralized water | 65 | 0.5556 | 0.4444 | 1.198 | 0.05095 | 0.2869 | 8.35 | 0.7379 | 75.11 |
| C_76 | demineralized water | 105 | 0.2972 | 0.7028 | 1.775 | 0.05621 | 0.3815 | 14.52 | 0.4817 | 127.6 |
| C_87 | demineralized water | 123 | 0.1523 | 0.8477 | 2.462 | 0.09534 | 0.5116 | 17.45 | 0.3021 | 139.2 |
| C_90 | demineralized water | 72 | 0.3935 | 0.6065 | 1.468 | 0.06026 | 0.3431 | 9.391 | 0.5712 | 96.43 |
| A_83 | 19.85 mM $^{15}NH_4Cl$ | 111 | 0.2567 | 0.7433 | 2.135 | 0.07621 | 0.4534 | 15.49 | 0.4694 | 165.1 |
| A_91 | 19.85 mM $^{15}NH_4Cl$ | 135 | 0.09471 | 0.9053 | 2.929 | 0.1385 | 0.597 | 19.46 | 0.2065 | 146.3 |
| A_99 | 19.85 mM $^{15}NH_4Cl$ | 118 | 0.5295 | 0.4705 | 1.454 | 0.03628 | 0.3048 | 16.63 | 0.7237 | 180 |
| A_103 | 19.85 mM $^{15}NH_4Cl$ | 55 | 0.4634 | 0.5366 | 1.375 | 0.0719 | 0.3431 | 6.896 | 0.6612 | 66.67 |
| N_79 | 19.98 mM $^{15}KNO_3$ | 83 | 0.1786 | 0.8214 | 2.128 | 0.1012 | 0.4816 | 11.06 | 0.312 | 99.87 |
| N_82 | 19.98 mM $^{15}KNO_3$ | 98 | 0.2927 | 0.7073 | 1.911 | 0.06899 | 0.4168 | 13.4 | 0.498 | 149.7 |
| N_94 | 19.98 mM $^{15}KNO_3$ | 105 | 0.3535 | 0.6465 | 1.793 | 0.05721 | 0.3853 | 14.52 | 0.5708 | 125.3 |
| N_102 | 19.98 mM $^{15}KNO_3$ | 126 | 0.1616 | 0.8384 | 2.495 | 0.0962 | 0.5159 | 17.95 | 0.3505 | 140.5 |

[a]Data was calculated from the rarefied OTU table at 97 % sequence similarity. The samples were rarified to 20051 reads per sample, where the number of reads is equal to the number of individuals per sample. Richness and diversity indices were calculated in the Paleontological Statistics Software (1).

**TABLE S2** List of KEGG pathways associated with the ectomycorrhizal fungi metatranscriptome[a]

| Nr. | Pathway map (number of matched objects) |
|---|---|
| 1 | lbc01100 Metabolic pathways - Laccaria bicolor (538) |
| 2 | lbc01110 Biosynthesis of secondary metabolites - Laccaria bicolor (245) |
| 3 | **lbc01230 Biosynthesis of amino acids - Laccaria bicolor (90)** |
| 4 | **lbc01200 Carbon metabolism - Laccaria bicolor (83)** |
| 5 | lbc00230 Purine metabolism - Laccaria bicolor (44) |
| 6 | lbc00520 Amino sugar and nucleotide sugar metabolism - Laccaria bicolor (43) |
| 7 | lbc00500 Starch and sucrose metabolism - Laccaria bicolor (37) |
| 8 | lbc00010 Glycolysis / Gluconeogenesis - Laccaria bicolor (35) |
| 9 | lbc00270 Cysteine and methionine metabolism - Laccaria bicolor (34) |
| 10 | lbc00620 Pyruvate metabolism - Laccaria bicolor (32) |
| 11 | lbc00564 Glycerophospholipid metabolism - Laccaria bicolor (31) |
| 12 | lbc00240 Pyrimidine metabolism - Laccaria bicolor (29) |
| 13 | **lbc00250 Alanine, aspartate and glutamate metabolism - Laccaria bicolor (29)** |
| 14 | lbc00970 Aminoacyl-tRNA biosynthesis - Laccaria bicolor (29) |
| 15 | lbc00561 Glycerolipid metabolism - Laccaria bicolor (28) |
| 16 | lbc00280 Valine, leucine and isoleucine degradation - Laccaria bicolor (27) |
| 17 | lbc00480 Glutathione metabolism - Laccaria bicolor (27) |
| 18 | lbc01210 2-Oxocarboxylic acid metabolism - Laccaria bicolor (27) |
| 19 | lbc00260 Glycine, serine and threonine metabolism - Laccaria bicolor (25) |
| 20 | lbc00020 Citrate cycle (TCA cycle) - Laccaria bicolor (24) |
| 21 | **lbc00330 Arginine and proline metabolism - Laccaria bicolor (23)** |
| 22 | lbc04146 Peroxisome - Laccaria bicolor (23) |
| 23 | lbc00030 Pentose phosphate pathway - Laccaria bicolor (22) |
| 24 | lbc00071 Fatty acid degradation - Laccaria bicolor (22) |
| 25 | lbc00380 Tryptophan metabolism - Laccaria bicolor (21) |
| 26 | lbc00562 Inositol phosphate metabolism - Laccaria bicolor (21) |
| 27 | lbc00630 Glyoxylate and dicarboxylate metabolism - Laccaria bicolor (21) |
| 28 | lbc00051 Fructose and mannose metabolism - Laccaria bicolor (18) |
| 29 | lbc00310 Lysine degradation - Laccaria bicolor (18) |
| 30 | lbc00770 Pantothenate and CoA biosynthesis - Laccaria bicolor (17) |
| 31 | **lbc00220 Arginine biosynthesis - Laccaria bicolor (16)** |
| 32 | lbc03050 Proteasome - Laccaria bicolor (16) |
| 33 | lbc01212 Fatty acid metabolism - Laccaria bicolor (15) |
| 34 | lbc03410 Base excision repair - Laccaria bicolor (15) |
| 35 | lbc04070 Phosphatidylinositol signaling system - Laccaria bicolor (15) |
| 36 | lbc00040 Pentose and glucuronate interconversions - Laccaria bicolor (14) |
| 37 | lbc00340 Histidine metabolism - Laccaria bicolor (14) |
| 38 | lbc00350 Tyrosine metabolism - Laccaria bicolor (14) |
| 39 | lbc00410 beta-Alanine metabolism - Laccaria bicolor (14) |
| 40 | lbc00680 Methane metabolism - Laccaria bicolor (14) |
| 41 | lbc00053 Ascorbate and aldarate metabolism - Laccaria bicolor (13) |
| 42 | lbc00640 Propanoate metabolism - Laccaria bicolor (13) |
| 43 | lbc03030 DNA replication - Laccaria bicolor (13) |
| 44 | lbc00400 Phenylalanine, tyrosine and tryptophan biosynthesis - Laccaria bicolor (12) |
| 45 | lbc00510 N-Glycan biosynthesis - Laccaria bicolor (12) |
| 46 | lbc00600 Sphingolipid metabolism - Laccaria bicolor (12) |
| 47 | lbc00650 Butanoate metabolism - Laccaria bicolor (12) |
| 48 | lbc00860 Porphyrin and chlorophyll metabolism - Laccaria bicolor (12) |
| 49 | lbc00920 Sulfur metabolism - Laccaria bicolor (12) |
| 50 | lbc04138 Autophagy - yeast - Laccaria bicolor (12) |
| 51 | lbc00300 Lysine biosynthesis - Laccaria bicolor (11) |
| 52 | lbc00760 Nicotinate and nicotinamide metabolism - Laccaria bicolor (11) |
| 53 | lbc00052 Galactose metabolism - Laccaria bicolor (10) |
| 54 | lbc00670 One carbon pool by folate - Laccaria bicolor (10) |
| 55 | lbc04011 MAPK signaling pathway - yeast - Laccaria bicolor (10) |
| 56 | lbc00740 Riboflavin metabolism - Laccaria bicolor (9) |
| 57 | lbc00130 Ubiquinone and other terpenoid-quinone biosynthesis - Laccaria bicolor (8) |

| | |
|---|---|
| 58 | lbc00360 Phenylalanine metabolism - Laccaria bicolor (8) |
| 59 | lbc00450 Selenocompound metabolism - Laccaria bicolor (8) |
| 60 | lbc00900 Terpenoid backbone biosynthesis - Laccaria bicolor (8) |
| 61 | **lbc00910 Nitrogen metabolism - Laccaria bicolor (8)** |
| 62 | lbc03008 Ribosome biogenesis in eukaryotes - Laccaria bicolor (8) |
| 63 | lbc03015 mRNA surveillance pathway - Laccaria bicolor (8) |
| 64 | lbc04113 Meiosis - yeast - Laccaria bicolor (8) |
| 65 | lbc00061 Fatty acid biosynthesis - Laccaria bicolor (7) |
| 66 | lbc00290 Valine, leucine and isoleucine biosynthesis - Laccaria bicolor (7) |
| 67 | lbc00430 Taurine and hypotaurine metabolism - Laccaria bicolor (7) |
| 68 | lbc00460 Cyanoamino acid metabolism - Laccaria bicolor (7) |
| 69 | lbc00513 Various types of N-glycan biosynthesis - Laccaria bicolor (7) |
| 70 | lbc00730 Thiamine metabolism - Laccaria bicolor (7) |
| 71 | lbc03013 RNA transport - Laccaria bicolor (7) |
| 72 | lbc03018 RNA degradation - Laccaria bicolor (7) |
| 73 | lbc04139 Mitophagy - yeast - Laccaria bicolor (7) |
| 74 | lbc00100 Steroid biosynthesis - Laccaria bicolor (6) |
| 75 | lbc00511 Other glycan degradation - Laccaria bicolor (6) |
| 76 | lbc00780 Biotin metabolism - Laccaria bicolor (6) |
| 77 | lbc00790 Folate biosynthesis - Laccaria bicolor (6) |
| 78 | lbc03420 Nucleotide excision repair - Laccaria bicolor (6) |
| 79 | lbc01040 Biosynthesis of unsaturated fatty acids - Laccaria bicolor (5) |
| 80 | lbc03020 RNA polymerase - Laccaria bicolor (5) |
| 81 | lbc04111 Cell cycle - yeast - Laccaria bicolor (5) |
| 82 | lbc04141 Protein processing in endoplasmic reticulum - Laccaria bicolor (5) |
| 83 | lbc04144 Endocytosis - Laccaria bicolor (5) |
| 84 | lbc00190 Oxidative phosphorylation - Laccaria bicolor (4) |
| 85 | lbc00261 Monobactam biosynthesis - Laccaria bicolor (4) |
| 86 | lbc00592 alpha-Linolenic acid metabolism - Laccaria bicolor (4) |
| 87 | lbc00750 Vitamin B6 metabolism - Laccaria bicolor (4) |
| 88 | lbc03450 Non-homologous end-joining - Laccaria bicolor (4) |
| 89 | lbc00062 Fatty acid elongation - Laccaria bicolor (3) |
| 90 | lbc00072 Synthesis and degradation of ketone bodies - Laccaria bicolor (3) |
| 91 | lbc00514 Other types of O-glycan biosynthesis - Laccaria bicolor (3) |
| 92 | lbc00515 Mannose type O-glycan biosynthesis - Laccaria bicolor (3) |
| 93 | lbc00565 Ether lipid metabolism - Laccaria bicolor (3) |
| 94 | lbc00590 Arachidonic acid metabolism - Laccaria bicolor (3) |
| 95 | lbc03040 Spliceosome - Laccaria bicolor (3) |
| 96 | lbc03430 Mismatch repair - Laccaria bicolor (3) |
| 97 | lbc00332 Carbapenem biosynthesis - Laccaria bicolor (2) |
| 98 | lbc00603 Glycosphingolipid biosynthesis - globo and isoglobo series - Laccaria bicolor (2) |
| 99 | lbc03060 Protein export - Laccaria bicolor (2) |
| 100 | lbc04136 Autophagy - other - Laccaria bicolor (2) |
| 101 | lbc00531 Glycosaminoglycan degradation - Laccaria bicolor (1) |
| 102 | lbc00563 Glycosylphosphatidylinositol (GPI)-anchor biosynthesis - Laccaria bicolor (1) |
| 103 | lbc00604 Glycosphingolipid biosynthesis - ganglio series - Laccaria bicolor (1) |
| 104 | lbc00660 C5-Branched dibasic acid metabolism - Laccaria bicolor (1) |
| 105 | lbc03440 Homologous recombination - Laccaria bicolor (1) |
| 106 | lbc04120 Ubiquitin mediated proteolysis - Laccaria bicolor (1) |
| 107 | lbc04122 Sulfur relay system - Laccaria bicolor (1) |
| 108 | lbc04145 Phagosome - Laccaria bicolor (1) |

[a]The mapping was performed in KEGG Mapper (1) using *Laccaria bicolor* as reference and 866 complete Enzyme Commission numbers assigned to the aggregated ectomycorrhizal fungi metatranscriptome.

**TABLE S3** Gene ontology significantly enriched molecular level functions and biological activities of *Fagus sylvatica* L DEGs in the nitrate treatment[a]

| GO Accession | Ontology | Description | FDR P value |
|---|---|---|---|
| GO:0015103 | MF | inorganic anion transmembrane transporter activity | 0.000475611 |
| GO:0015112 | MF | nitrate transmembrane transporter activity | 0.001682898 |
| GO:0015318 | MF | inorganic molecular entity transmembrane transporter activity | 0.02315464 |
| GO:0008509 | MF | anion transmembrane transporter activity | 0.02315464 |
| GO:0015075 | MF | ion transmembrane transporter activity | 0.02315464 |
| GO:0008308 | MF | voltage-gated anion channel activity | 0.025644629 |
| GO:0005216 | MF | ion channel activity | 0.031224718 |
| GO:0005253 | MF | anion channel activity | 0.031224718 |
| GO:0005244 | MF | voltage-gated ion channel activity | 0.032827156 |
| GO:0016710 | MF | trans-cinnamate 4-monooxygenase activity | 0.032827156 |
| GO:0019150 | MF | D-ribulokinase activity | 0.032827156 |
| GO:0042300 | MF | beta-amyrin synthase activity | 0.032827156 |
| GO:0009671 | MF | nitrate:proton symporter activity | 0.032827156 |
| GO:0008909 | MF | isochorismate synthase activity | 0.032827156 |
| GO:0016712 | MF | oxidoreductase activity, acting on paired donors, with incorporation or reduction of molecular oxygen, reduced flavin or flavoprotein as one donor, and incorporation of one atom of oxygen | 0.032827156 |
| GO:0015267 | MF | channel activity | 0.032827156 |
| GO:0022832 | MF | voltage-gated channel activity | 0.032827156 |
| GO:0022803 | MF | passive transmembrane transporter activity | 0.032827156 |
| GO:0062047 | MF | pipecolic acid N-hydroxylase | 0.032827156 |
| GO:0098809 | MF | nitrite reductase activity | 0.032827156 |
| GO:0017096 | MF | acetylserotonin O-methyltransferase activity | 0.032827156 |
| GO:0016662 | MF | oxidoreductase activity, acting on other nitrogenous compounds as donors, cytochrome as acceptor | 0.032827156 |
| GO:0050421 | MF | nitrite reductase (NO-forming) activity | 0.032827156 |
| GO:0019825 | MF | oxygen binding | 0.041848523 |
| GO:0016211 | MF | ammonia ligase activity | 0.041848523 |
| GO:0080019 | MF | fatty-acyl-CoA reductase (alcohol-forming) activity | 0.041848523 |
| GO:0015296 | MF | anion:cation symporter activity | 0.041848523 |
| GO:0004356 | MF | glutamate-ammonia ligase activity | 0.041848523 |

| | | | |
|---|---|---|---|
| GO:0015098 | MF | molybdate ion transmembrane transporter activity | 0.041848523 |
| GO:0031559 | MF | oxidosqualene cyclase activity | 0.041848523 |
| GO:0015513 | MF | high-affinity secondary active nitrite transmembrane transporter activity | 0.041848523 |
| GO:0004016 | MF | adenylate cyclase activity | 0.041848523 |
| GO:0004820 | MF | glycine-tRNA ligase activity | 0.041848523 |
| GO:0004345 | MF | glucose-6-phosphate dehydrogenase activity | 0.041848523 |
| GO:0050486 | MF | intramolecular transferase activity, transferring hydroxy groups | 0.041848523 |
| GO:0022857 | MF | transmembrane transporter activity | 0.041848523 |
| GO:0022836 | MF | gated channel activity | 0.045553402 |
| GO:0016866 | MF | intramolecular transferase activity | 0.046986661 |
| GO:0015698 | BP | inorganic anion transport | 0.000685462 |
| GO:0010167 | BP | response to nitrate | 0.003443921 |
| GO:0015706 | BP | nitrate transport | 0.003443921 |
| GO:0006821 | BP | chloride transport | 0.03664734 |

[a]Analysis carried out in g:Profiler (Version: e101_eg48_p14_baf17f0) with *Arabidopsis thaliana* as reference; statistical domain scope: custom over annotated genes; significance threshold: FDR P-adjusted < 0.05).

**TABLE S4** Mean ± standard deviation values of air temperature and humidity for the duration of the fertilization experiment and monthly averages from date of tree collection to the date of harvest

| Date | Air temperature (°C) | Air humidity (%) |
|---|---|---|
| 17 July 2018 to 19 July 2018 | 22.8 ± 4.47 | 52.57 ± 8.32 |
| 31 July 2018 to 2 August 2018 | 27.16 ± 3.58 | 54.70 ± 11.77 |
| March | 4.72 ± 5.00 | 78.34 ± 16.04 |
| April | 14.16 ± 5.86 | 68.10 ± 18.53 |
| May | 17.74 ± 6.05 | 64.45 ± 19.12 |
| June | 18.87 ± 5.16 | 70.70 ± 17.76 |
| July | 22.40 ± 6.46 | 57.99 ± 20.59 |
| August | 26.34 ± 4.70 | 57.83 ± 14.81 |

**TABLE S5** Overview of root-associated fungi sequence processing results based on Illumina MiSeq of the fungal ITS2 rRNA gene. Fungal reads were rarefied to 20051 reads, the lowest sequencing depth among samples

| Sample | Treatment | Raw reads | Filtered reads | Processed reads (%) | Number of fungal reads | Number of fungal OTUs before rarefaction | Number of fungal OTUs after rarefaction |
|---|---|---|---|---|---|---|---|
| C_74 | demineralized water | 81455 | 76478 | 93.89 | 47733 | 85 | 65 |
| C_76 | demineralized water | 33591 | 31327 | 93.26 | **20051** | 105 | 105 |
| C_87 | demineralized water | 65622 | 61057 | 93.04 | 32810 | 132 | 123 |
| C_90 | demineralized water | 45123 | 41405 | 91.76 | 26636 | 75 | 72 |
| A_83 | 19.85 mM $^{15}NH_4Cl$ | 57103 | 53497 | 93.69 | 41015 | 127 | 111 |
| A_91 | 19.85 mM $^{15}NH_4Cl$ | 86563 | 80679 | 93.20 | 50894 | 159 | 135 |
| A_99 | 19.85 mM $^{15}NH_4Cl$ | 60363 | 56470 | 93.55 | 32104 | 126 | 118 |
| A_103 | 19.85 mM $^{15}NH_4Cl$ | 45808 | 43041 | 93.96 | 33304 | 72 | 55 |
| N_79 | 19.98 mM $^{15}KNO_3$ | 72175 | 67093 | 92.96 | 40852 | 104 | 83 |
| N_82 | 19.98 mM $^{15}KNO_3$ | 48344 | 45565 | 94.25 | 35446 | 111 | 98 |
| N_94 | 19.98 mM $^{15}KNO_3$ | 49988 | 46847 | 93.72 | 31601 | 120 | 105 |
| N_102 | 19.98 mM $^{15}KNO_3$ | 43122 | 40659 | 94.29 | 31177 | 139 | 126 |

**TABLE S6** Overview of RNA sequence processing and mapping statistics

| Sample | Treatment | RNA Integrity Number | Raw reads | Processed reads | % processed reads | Mapped reads | Mapped to fungi | Mapped to Fagus | % mapped | % mapped fungi | % mapped Fagus |
|---|---|---|---|---|---|---|---|---|---|---|---|
| C_74 | demineralized water | 6.7 | 117,404,490 | 116,514,831 | 99.24 | 75,838,597 | 38,345,820 | 37,492,777 | 65.09 | 32.91 | 32.18 |
| C_76 | demineralized water | 7.0 | 110,533,021 | 109,713,595 | 99.26 | 64,762,941 | 5,678,950 | 59,083,991 | 59.03 | 5.18 | 53.85 |
| C_87 | demineralized water | 7.9 | 123,830,699 | 123,001,454 | 99.33 | 78,074,569 | 3,662,184 | 74,412,385 | 63.47 | 2.98 | 60.5 |
| C_90 | demineralized water | 7.6 | 113,626,927 | 112,848,395 | 99.31 | 73,581,581 | 33,664,910 | 39,916,671 | 65.2 | 29.83 | 35.37 |
| A_83 | 19.85 mM $^{15}NH_4Cl$ | 7.6 | 113,949,314 | 113,135,195 | 99.29 | 68,432,056 | 29,792,675 | 38,639,381 | 60.49 | 26.33 | 34.15 |
| A_91 | 19.85 mM $^{15}NH_4Cl$ | 7.4 | 101,039,883 | 100,271,972 | 99.24 | 56,633,842 | 3,005,149 | 53,628,693 | 56.48 | 3 | 53.48 |
| A_99 | 19.85 mM $^{15}NH_4Cl$ | 7.1 | 111,430,088 | 110,558,237 | 99.22 | 70,563,179 | 18,636,615 | 51,926,564 | 63.82 | 16.86 | 46.97 |
| A_103 | 19.85 mM $^{15}NH_4Cl$ | 7.5 | 109,734,463 | 108,965,024 | 99.3 | 73,756,153 | 23,338,402 | 50,417,751 | 67.69 | 21.42 | 46.27 |
| N_79 | 19.98 mM $^{15}KNO_3$ | 7.3 | 102,541,237 | 101,839,654 | 99.32 | 62,315,939 | 11,130,991 | 51,184,948 | 61.19 | 10.93 | 50.26 |
| N_82 | 19.98 mM $^{15}KNO_3$ | 7.0 | 98,286,285 | 97,604,837 | 99.31 | 57,297,015 | 26,933,739 | 30,363,276 | 58.7 | 27.59 | 31.11 |
| N_94 | 19.98 mM $^{15}KNO_3$ | 7.8 | 113,990,445 | 113,207,752 | 99.31 | 66,413,204 | 5,254,953 | 61,158,251 | 58.66 | 4.64 | 54.02 |
| N_102 | 19.98 mM $^{15}KNO_3$ | 7.1 | 107,765,136 | 107,032,242 | 99.32 | 57,103,575 | 13,922,719 | 43,180,856 | 53.35 | 13.01 | 40.34 |

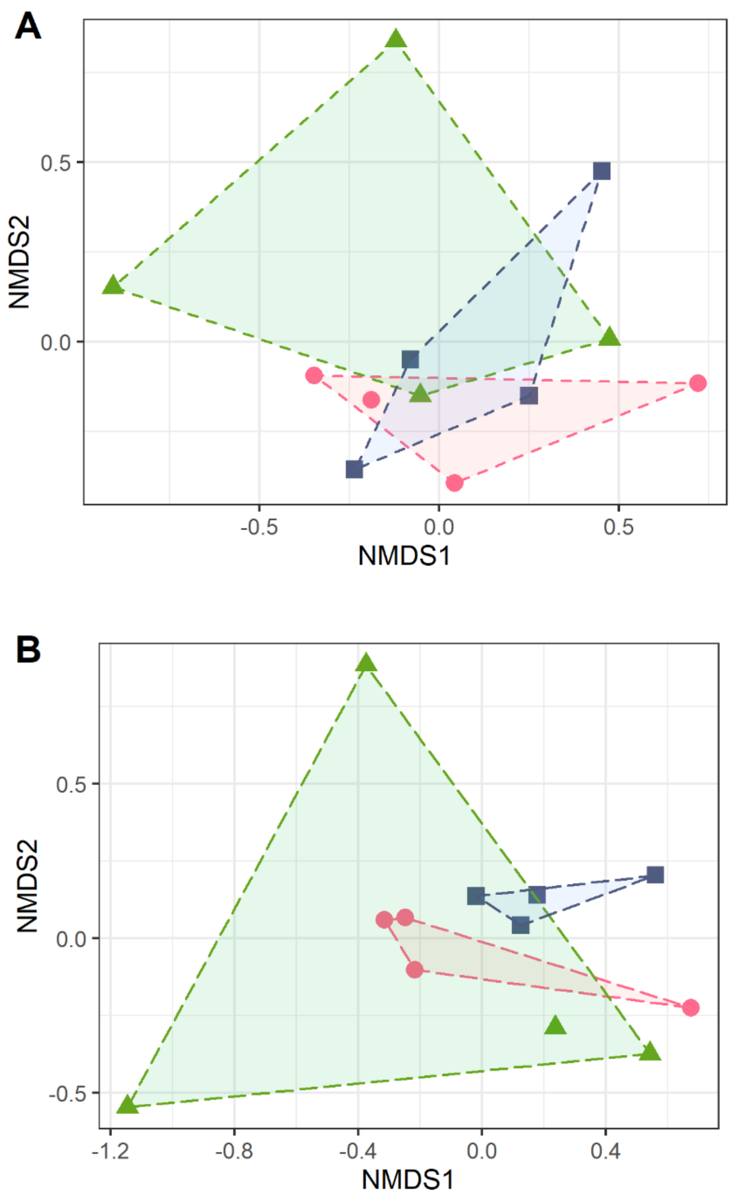


**FIG S1** Non-metric multidimensional scaling (NMDS) ordination of the fungal community structure at the OTU level (A), and NMDS ordination of the fungal raw RNA counts characterized by taxonomy (B) in control (triangle), ammonium (circle) and nitrate (square) treated beech root tips. Ordinations are based on Hellinger-transformed data with Bray-Curtis dissimilarity. A: stress = 0.12; there was no significant difference in the fungal community composition among treatment conditions ($R^2$= 0.146, pseudo-$F_{2,9}$= 0.767, p= 0.861, permutations = 9999). B: stress = 0.08; there was no significant difference in the fungal metatranscriptomes among treatment conditions ($R^2$= 0.198, pseudo-$F_{2,9}$= 1.110, p= 0.353, permutations = 9999, adonis).

**TEXT S1** Methodological details

***Ammonium and nitrate measurements from soil.*** Before the $^{15}N$ treatments, ammonium was extracted from soil using 1mM CaCl2 solution, the extracts were freezed-dried and re-solved in 0.5 mL ultrapure water, then a 1:10 diluted aliquot was used for photometric measurements at 690 nm (Beckmann Spectrophotometer DU-640) using the ammonium test kit 100683 and ammonium standard solution 119812 from Merck Millipore (Merck KGaA, Darmstadt, Germany), according to the manufacturer instructions. Nitrate was also extracted from soil using 1mM CaCl2 solution, the extracts were freeze-dried, re-solved in 0.5 mL ultrapure water, and then a 1:10 diluted aliquot was used for photometric measurements at 340 nm (Beckmann Spectrophotometer DU-640) using the nitrate test kit 109713 and nitrate standard solution 119811 from Merck Millipore (Merck KGaA, Darmstadt, Germany), according to the manufacturer instructions.

**Ammonium and nitrate measurements in fine roots.** Ammonium and nitrate were extracted from fine roots after (1) with slight modifications. In detail, fresh fine roots, previously shock-frozen in liquid nitrogen and stored at - 80 °C during harvest time, were milled under liquid nitrogen using a ball mill Type MM400 (Retsch GmbH, Haan, Germany) in stainless steel grinding jars at a frequency of 30/s in intervals of 20 s to avoid thawing. For nitrate extraction, approximately 55 mg (exact weight recorded) of frozen milled sample were placed inside a 2 mL screw cap micro tube (Order number: 72.693.100, Sarstedt AG & Co. KG, Nümbrecht, Germany) with 500 µL of pre-heated (80 °C) autoclaved ultrapure water (arium® pro, Water Purification System, Sartorius Lab Instruments GmbH & Co. KG, Goettingen, Germany) to denature nitrate reductase. The sample was incubated with the lid closed at 100 °C and 2000 rpm for 20 min in an Eppendorf ThermoMixer C (model: 5382, Eppendorf AG, Hamburg, Germany). The sample was cooled on ice and centrifuged at 20,400 x g (Eppendorf centrifuge 5427 R, Eppendorf AG, Hamburg, Germany) for 10 min at room temperature. The supernatant was aliquoted, shock-frozen in liquid nitrogen and stored at - 80 °C. For ammonium extraction, approximately 55 mg (exact weight recorded) of frozen milled sample were placed inside a 2 mL screw cap micro tube (Sarstedt AG & Co. KG) on ice with 1000 µL of 0.1 M hydrochloric acid (Catalogue No. 100316, Merck KGaA, Darmstadt, Germany) prepared with autoclaved ultrapure water (Sartorius Lab Instruments GmbH & Co. KG). The sample was mixed gently for 10 s until a homogenous solution was formed. The sample was place on ice and 500 µL of chloroform (Art. No. 4432.1, Carl Roth GmbH & Co. KG, Karlsruhe, Germany) were added. The sample was incubated with the lid closed at 4 °C and 2000 rpm for 15 min the thermomixer. The extract was centrifuged at 12,000 x g (Eppendorf 5427 R) at 8 °C for 10 minutes. The top aqueous phase was transferred into a new 1.5 mL reaction tube containing 50 mg of acid-washed activated carbon from peat (Labochem international, neoFroxx GmbH, Einhausen, Germany, distributed by NeoLab Migge GmbH, product number: LC-10810.3) on ice. The sample was mixed for 10 s to form a homogenous solution. The sample was centrifuged at 20,400 x g (Eppendorf 5427 R) at 8 °C for 5 min and the supernatant was transferred into a new reaction tube on ice. The sample was centrifuged again at 20,400 x g (Eppendorf 5427 R) at 8 °C for 5 min and the supernatant was transferred into a new reaction tube on ice. Sedimentation of free-floating light weight acid-washed carbon particles in the sample was induced by shock-freezing the sample in liquid nitrogen and thawing on ice. The sample was centrifuged at 20,4000 x g (Eppendorf 5427 R) at 8 °C for 5 min and the supernatant was transferred to a new tube on ice. The extract (transparent in color) was aliquoted, shock-frozen in liquid nitrogen and stored at - 80 °C. Nitrate and ammonium were measured spectrophotometrically in triplicates with commercially available kits (Spectroquant® Nitrate Test Kit Merk, KGaA, Darmstadt, Germany, catalog number: 1.09713.0002; Spectroquant® Ammonium Test Kit, Merk, KGaA, Darmstadt, Germany, catalog number: 1.14752.0002) according to the guidelines of the manufacturer. Reagent quantities were adjusted to the volume of the reaction (i.e., for nitrate: 100 µL of sample, 800 µL of reagent 1 and 100 µL of reagent 2; for ammonium: 700 µL of sample, 84 µL of reagent 1, 16 mg of reagent 2 and 11.2 µL of reagent 3). Autoclaved ultrapure water (Sartorius Lab Instruments GmbH & Co. KG) was used instead of root extract as blank. Extracts were diluted when necessary, with autoclaved ultrapure water. When the reaction was accomplished, the sample was transferred into a disposable cuvette (ammonium: Semi-micro cuvette, Ref: 67.742 Sarstedt AG & Co. KG, Nümbrecht, Germany; nitrate: UV-cuvette semi-micro, Cat. No. 759150, Brand GmbH & Co. KG, Wertheim, Germany) and light absorbance was measured at 690 nm for ammonium and 340 nm for nitrate using a spectrophotometer (model: DU640, Beckman Instruments, Inc., Fullerton, California, USA) against the blank reaction as background. Standard curves were prepared in the range of the detection limits of the reaction kits and used for the determination of the nitrate and ammonium concentrations in the extracts.

**Non-structural carbohydrate measurements in tree fine roots.** Glucose, fructose, sucrose and starch concentrations in fine root extracts were measured enzymatically in three technical replicates per sample by measuring the generation of NADPH (2). Fine root aliquots from -80°C were milled in liquid nitrogen using a ball mill Type MM400

(Retsch GmbH, Haan, Germany). Approximately 75 mg of milled sample (exact weight recorded) were placed in a 2 mL screw cap micro tube (Order number: 72.693.100, Sarstedt AG & Co. KG, Nümbrecht, Germany) and extracted with 1500 µL of extraction medium (pH 0.45) containing 80 : 20 (v : v) dimethyl sulphoxide : 25% hydrochloric acid (Art. No. 4720.2, Carl Roth GmbH, Karlsruhe, Germany; Catalogue No. 100316, Merck KGaA, Darmstadt, Germany). The sample was mixed for 10 seconds, incubated at 60 °C at 2000 rpm for 30 min in an Eppendorf ThermoMixer C (model: 5382, Eppendorf AG, Hamburg, Germany) and then cooled on ice. The sample was centrifuged at 5000 x g for 10 min at 4 °C (Eppendorf 5427 R, Eppendorf AG, Hamburg, Germany) and the supernatant was transferred to a new reaction tube on an ice bath. A 200 µL aliquot of the extract was adjusted to pH 4.6 with 1200 µL of cold 0.2 M citrate buffer (pH 10.6) (Product number: S4641, Sigma-Aldrich Chemie GmbH, Taufkirchen, Germany). For glucose, fructose and sucrose measurements, 1 volume (400 µL) of the 7x diluted (pH 4.6) extract was mixed with 1 volume (400 µL) of cold 50 µM citrate buffer (pH 4.6) (Sigma-Aldrich Chemie GmbH) to make a 14x diluted sample at pH 4.6. Glucose and fructose were measured in the same cuvette by consecutive addition of enzymes. For this purpose, a 100 µL aliquot of the 14x diluted sample was mixed in a disposable UV-cuvette semi-micro (Cat. No. 759150, Brand GmbH & Co. KG, Wertheim, Germany) with 400 µL of autoclaved ultrapure water (arium® pro, Water Purification System, Sartorius Lab Instruments GmbH & Co. KG, Göttingen, Germany) and 250 µL of coenzyme solution (pH 7.6), which was prepared in 0.75 M triethanolamine buffer (Catalogue No. 1.08379, Merck KGaA, Darmstadt, Germany) adjusted to pH 7.6 with 25% hydrochloric acid (Catalogue No. 100316, Merck KGaA, Darmstadt, Germany). The coenzyme solution contained 4mM

NADP (NADP-RO, Roche Diagnostics GmbH, Mannheim, Germany, product code: 10128058001, distributed by Merck KGaA, Darmstadt, Germany), 10 mM ATP (ATPD-RO, Roche Diagnostics GmbH, Mannheim, Germany, product code: 10127531001, distributed by Merck KGaA, Darmstadt, Germany) and 9 mM $MgSO_4$ (Art. No. P027.2, Carl Roth GmbH, Karlsruhe, Germany). After mixing with a stirring rod (Ref: 81.970, Sarstedt AG & Co. KG, Nümbrecht, Germany) and incubating for 3 min at 25 °C in the dark, the background absorbance of NADPH was measured at 340 nm at 25 °C using a spectrophotometer (model: DU640, Beckman Instruments, Inc., Fullerton, California, USA) and air as reference. To measure glucose, 10 µL of enzyme hexokinase/glucose-6-phosphate dehydrogenase (HK/G6P-DH-RO, 3 mg $mL^{-1}$, 340 U hexokinase $mL^{-1}$ and 170 U glucose-6-phosphate dehydrogenase $mL^{-1}$ at +25 °C, Roche Diagnostics GmbH, Mannheim, Germany, product code: 10737275001, distributed by Merck KGaA, Darmstadt, Germany) were added to the cuvette, mixed and incubated for 5 min at 25 °C in the dark before measuring the absorbance at 340 nm at 25 °C. For fructose measurement, 5 µL of enzyme phosphoglucose isomerase (PGI-RO, 10 mg $mL^{-1}$, ~350 units $mg^{-1}$ protein at 25 °C, Roche Diagnostics GmbH, Mannheim, Germany, product code: 10128139001, distributed by Merck KGaA, Darmstadt, Germany) were added to the same cuvette, mixed and incubated for 5 min in the dark before measuring the absorbance at 340 nm at 25 °C. The absorbance of glucose was subtracted from the absorbance of the fructose assay prior to calculating the fructose concentration. For sucrose, 100 µL aliquots of the 14x diluted extract were incubated for 15 min at 25 °C in a new cuvette with 10 µL of pre-warmed (37 °C) enzyme invertase solution (60 mg $mL^{-1}$, 200-300 U $mg^{-1}$ at 25 °C, Ref: I9274, Sigma-Aldrich Chemie GmbH, Taufkirchen, Germany), which was dissolved in 0.32 M citrate buffer at pH 4.6 (Sigma-

Aldrich Chemie GmbH). Next, 400 µL of autoclaved ultrapure water (arium® pro) and 250 µL of the coenzyme solution (pH 7.6) were added. After mixing and incubating for 3 min at 25 °C in the dark, the background absorbance of NADPH was measured at 340 nm with air as a reference. Then, 10 µL of the enzyme hexokinase/glucose-6-phosphate dehydrogenase (Roche Diagnostics GmbH) were added to the cuvette, mixed and incubated for 15 min at 25 °C in the dark before measuring the new absorbance of NADPH at 340 nm at 25 °C. The absorbance of free glucose, previously measured in the glucose assay of each sample, was subtracted from the absorbance of the sucrose assay for sucrose calculations. For starch measurements, an aliquot of 200 µL of the 7x diluted sample extract (pH 4.6) was mixed in a reaction vial with 200 µL enzyme amyloglucosidase solution (~70 U $mg^{-1}$, 2.64 mg $mL^{-1}$, product number: 10115, Sigma-Aldrich Chemie GmbH, Taufkirchen, Germany), which was dissolved in in 50 µM citrate buffer at pH 4.6 (Sigma-Aldrich Chemie GmbH). The sample was mixed, incubated at 57 °C (Thermo block type: 51336101, Gebr. Liebisch GmbH & Co. KG, Bielefeld, Germany) for 20 minutes and then cooled on ice. An aliquot of 100 µL of this sample was mixed with 400 µL of autoclaved ultrapure water (arium® pro) and 250 µL of the coenzyme solution. After mixing and incubating for 3 min at 25 °C, the background absorbance of NADPH was measured at 340 nm. Then, 10 µL of the enzyme hexokinase/glucose-6-phosphate dehydrogenase (Roche Diagnostics GmbH) were added, mixed and incubated for 5 min at 25 °C before measuring the absorbance at 340 nm at 25 °C. The absorbance of free glucose, previously measured in each sample in the glucose assay, was subtracted from the absorbance of the starch assay before starch calculations. All non-structural carbohydrates were measured in triplicates. Blank and standard reactions were run along in triplicates per batch of eight samples. The absorbance of the blank was subtracted from the absorbance of the respective carbohydrate's assay before subtraction of free glucose. The carbohydrate concentrations were calculated using the extinction coefficient ε = 6.3 (L x $mmol^{-1}$ x $cm^{-1}$) of NADPH at 340 nm and taking the dilution of the sample into account.

**Nucleic acid extractions, library preparation and sequencing.** Root apices stored at - 80 °C were homogenized in liquid nitrogen using sterilized mortar and pestle (180 °C, 4 hours). The powdered frozen sample was split in two parts: one for DNA extraction and Illumina sequencing of fungi and the other for RNA extraction and sequencing. **DNA extraction and Illumina sequencing of fungi associated to beech roots.** DNA was extracted from approximately 200 mg homogenized root apices using the innuPREP Plant DNA Kit (Analytik Jena, AG, Jena, Germany). The extracted DNA was purified with the DNeasy PowerClean Pro CleanUp Kit (Qiagen, Hilden, Germany). The concentration of the purified DNA was measured with a Qubit™ 3.0 Fluorometer using the Qubit™ dsDNA HS Assay Kit (Life Technologies GmbH, Darmstadt, Germany). The fungal nuclear ribosomal internal transcribed spacer (ITS2) region was amplified by Polymerase Chain Reaction (PCR) using the primer pair ITS3_KYO2 (3) and ITS4 (4), both containing specific Illumina overhang adapters as follows:

forward (Miseq_ITS4_KYO2): *TCGTCGGCAGCGTCAGATGTGTATAAGAGACAG*GATGAAGAACGYAGYRAA and reverse (Miseq_ITS4): *GTCTCGTGGGCTCGGAGATGTGTATAAGAGACAG*TCCTCCGCTTATTGATATGC with the adapters in italics and the primers underlined.

Triplicate PCR reactions were carried out per sample in independent runs and randomized order. Each reaction contained 13.65 µL of master mix: 1X Phusion HF buffer (Catalog No. F-530XL, Thermo Fisher Scientific Baltics UAB, Vilnius, Lithuania), 1.65 mM $MgCl_2$ (Catalog No. F-530XL, Thermo Fisher Scientific Baltics UAB, Vilnius, Lithuania), 200 µM of each dNTP (Catalog No. R0181, Thermo Fisher Scientific Baltics UAB, Vilnius, Lithuania), 0.2 µM forward primer with Illumina overhang adapter (Microsynth AG, Balgach, Switzerland), 0.2 µM reverse primer with Illumina overhang adapter (Microsynth AG) and 1 U of Phusion HF DNA polymerase (Catalog No. F-530XL, Thermo Fisher Scientific Baltics UAB, Vilnius, Lithuania). To each reaction, 150 ng of genomic DNA template was added, and the reaction was adjusted to a final volume of 50 µL with nuclease free water (Order No. A7398, AppliChem GmbH, Darmstadt, Germany. One positive control containing fungal DNA as template and one negative control containing nuclease free water instead of DNA as template were run per batch. The PCR reaction was carried out in a SensoQuest Labcycler (SensoQuest GmbH, Goettingen, Germany) with the following conditions: initial template denaturation at 98 °C for 2 min, 25 cycles of a second denaturation step at 98 °C for 10 s, annealing of primers at 48 °C for 20 s and elongation at 72 °C for 20 s, then a final elongation step at 72 °C for 5 min.

The PCR amplicons were checked by agarose gel (2.5 %) electrophoresis (Art No. 840004, Biozym Scientific GmbH, Hessisch Oldendorf, Germany). The GeneRuler 100 bp Plus DNA ladder (Catalog No. SM0322, Thermo Fisher Scientific Baltics UAB, Vilnius, Lithuania) was used for controlling the amplicon size. In-gel staining was done with ethidium bromide at a concentration of 0. 5 µg/mL (Carl Roth, Art Nr. 2218.1). Gel electrophoresis was done in 1xTAE buffer for 30 min at 80 volts. The gel was visualized under UV light and documented in a Gel IX Imager Gel Documentation System (Intas Science Imaging Instruments GmbH, Göttingen, Germany). The PCR amplicons were purified with magnetic beads using the Left Side Size Selection protocol of the Magsi-NGSPREP Plus kit (Steinbrenner Laborsysteme GmbH, Wiesenbach, Germany) according to the manufacturer's instructions. The concentration of purified PCR amplicons was measured with the Qubit™ 3.0 fluorometer using the Qubit™ dsDNA HS Assay Kit. The three technical PCR replicates per sample were equimolarly pooled in nuclease free reaction tubes. The concentration of the pooled PCR amplicons was measured once again with the Qubit™ 3.0 fluorometer using the Qubit™ dsDNA HS Assay Kit. Finally, the pooled PCR amplicons were delivered to the Goettingen Genomics Laboratory (G2L, Institute of Microbiology and Genetics, University of Goettingen, Goettingen, Germany) where an indexing PCR was carried out using eight additional PCR cycles prior to sequencing. The indexing PCR step was done to ligate unique Illumina identifiers (a.k.a., indexes, barcodes, tags) to the adapters already attached to the amplicons of each sample. This allows for simultaneous sequencing of amplicons from different samples in one run and separation of reads according to sample after sequencing. After indexing PCR, amplicon DNA concentration was quantified with the Qubit™ 3.0 Fluorometer (LifeTechnologies GmbH), then amplicons were pooled at equimolar concentration and sequenced on a MiSeq flow cell using Reagent Kit v3 and 2x300 pair-end reads (Illumina Inc., San Diego, USA) according to the manufacturer's instructions.

**Fungal ITS2 amplicon sequence bioinformatic processing and data analyses.** Raw pair-end read quality filtering was performed with fastp v0.20.0 (5). Reads with an average quality score lower than 20 were removed, sequences were clipped via sliding window of 4 (minimum phredscore of 20) and corrected by overlap. Pair-end reads were merged with PEAR v0.9.11 (6). Forward and reverse primers were removed using cutadapt 2.5 with Python 3.7.3. The tool VSEARCH (7) v2.14.1 was used for size filtering (sequences shorter than 140 bp were removed), dereplication (100 % threshold), denoised (UNOISE3), chimeric sequence removal (UCHIME3, de novo and reference based), sorting sequences by cluster size and clustering of OTUs at 97% sequence identity. The merged pair-end reads were mapped to operational taxonomic units (OTUs) and abundance tables were generated. Taxonomic assignment of OTUs was carried out against the UNITE database v8.2 (04.02.2020) (8). All unidentified ASVs were searched (blastn) (9) against the nt database (2020-01-17) to remove non-fungal ASVs and only fungal sequence reads were kept. Finally, extrinsic domain ASVs and unclassified ASVs were discarded from the taxonomic table. The fungal OTUs were assigned into ecological trophic modes for identification of functional groups using the FUNGUild annotation tool (10). The sequencing depth was controlled by rarefaction analysis and the samples were normalized by rarefying to the sample having the lowest sequencing depth (i.e. 20051 sequence reads) using the package ampvis2 (11). Because all non-fungal sequences were removed, sequences classified as “unknown eukaryotes” in the OTU table represent fungi with unknown phylogenetic lineage.

**RNA extraction from beech root apices, library preparation and sequencing of beech and associated fungi.** Total RNA was isolated from beech root apices using a modified method after (12). In detail, frozen root apices (-80°C) were homogenized to a fine powder using sterilized (180 °C, 4 h) mortar and pestle and liquid nitrogen. Approximately 150 mg of frozen homogenized sample were placed inside a 2 mL micro tube (Ref: 72.695.500, Sarstedt AG & Co. KG, Nümbrecht, Germany) and extracted with 800 µL of pre-heated (65 °C) extraction buffer containing: 2% (w/v) hexadecyltrimethylammonium bromide (CTAB, Art. No. 9161.1, Carl Roth GmbH & Co. KG, Karlsruhe, Germany), 2% (w/v) polyvinylpyrrolidinone K 30 (PVP, Art. No. 4607.2, Carl Roth GmbH & Co. KG, Karlsruhe, Germany), 100 mM Tris HCl (pH 8.0) (Art. No: 9090.2, Carl Roth GmbH & Co. KG, Karlsruhe, Germany), 25 mM EDTA (Product number: A5097,0500, AppliChem GmbH, Darmstadt, Germany) and 2.0 M NaCl (Art. No: 3957.1, Carl Roth GmbH & Co. KG, Karlsruhe, Germany). The extraction buffer was prepared in ultrapure water (arium® pro, Water Purification System, Sartorius Lab Instruments GmbH & Co. KG, Goettingen, Germany), treated with 0.1 % diethyl pyrocarbonate (D5758, Sigma-Aldrich, Chemie GmbH, Steinheim, Germany) and filter-sterilized (0.22 µM) (Ref: 83.1826.001, Sarstedt AG & Co. KG, Nümbrecht, Germany). Immediately, 16 µL of 2-mercaptoethanol (4 °C) (Art. No: 4227.3, Carl Roth GmbH & Co. KG, Karlsruhe, Germany) were added to each sample. The sample was mixed for 10 s until a homogenous suspension was formed. The sample was incubated for 15 min at 65 °C at 1000 rpm (Eppendorf ThermoMixer® C, model: 5382, Eppendorf AG, Hamburg, Germany). The sample was incubated at room temperature for approximately 15 min while being regularly inverted horizontally. Then, 800 µL of chloroform:isoamyl alcohol (24:1) (v:v) (Art. No. 4432.1, Carl Roth

GmbH & Co. KG, Karlsruhe, Germany; product number: 1.00979, Merck KGaA, Darmstadt, Germany) were added to the CTAB extraction mixture and the sample was mixed for 3 s. The sample was incubated in for 15 min at 22 °C at 1400 rpm in the thermomixer. To separate the phases, the sample was centrifuged at 20800 x g (Eppendorf centrifuge 5417 R, Eppendorf AG, Hamburg, Germany) at room temperature °C for 5 min. The upper phase was transferred into a new 2 mL micro tube avoiding flocculent material at the interface. The lower phase and interphase were discarded in the appropriate chloroform waste. Another 800 µL of chloroform:isoamyl alcohol (24:1) (v:v) (Carl Roth GmbH & Co. KG; Merck KGaA) were added to the new 2 mL tube containing the upper phase and the sample was vortexed for 3 s. The sample was centrifuged at 20800 x g (Eppendorf 5417 R) at room temperature for 5 min to separate the phases. The upper phase was transferred again into a new 2 mL micro tube avoiding flocculent material at the interface. The lower phase and interphase were discarded in the appropriate chloroform waste. The washing steps with 800 µL of chloroform:isoamyl alcohol (24:1) (v:v) (Carl Roth GmbH & Co. KG; Merck KGaA) were repeated two additional times or until visible flocculent material (proteins, lipids and carbohydrates) lingering within the aqueous layer near the interface were no longer visible in the supernatant. After repeated washing steps, the upper phase was finally transferred into a 1.5 mL SafeSeal tube (Ref: 72.706, Sarstedt AG & Co. KG) on ice. One fourth times the sample volume (i.e., 0.25 x 180 µL to 200 µL = 45 µL to 50 µL) of cold (-20 °C) 10 M LiCl (Art. No. 6698.2, Carl Roth GmbH & Co. KG, Karlsruhe, Germany) were added to the upper phase in the 1.5 mL tube. The sample was mixed for 3 s. The RNA precipitated overnight on ice at 4 °C. After 12 h, the sample was centrifuged at 20800 x g (Eppendorf 5417 R) at 4 °C for 20 min. The upper phase was carefully discarded

(CTAB waste). Then, 400 µL of preheated (65 °C) SSTE buffer (pH 8.0) containing: 0.5% (w/v) SDS ultra (Art. No: 2326.2, Carl Roth GmbH & Co. KG, Karlsruhe, Germany), 10 mM Tris-HCl (Art. No: 9090.2, Carl Roth GmbH & Co. KG, Karlsruhe, Germany), 1 mM EDTA (Product number: A5097,0500, AppliChem GmbH, Darmstadt, Germany) and 1 M NaCl (Art. No: 3957.1, Carl Roth GmbH & Co. KG, Karlsruhe, Germany) were added to the pellet. The SSTE buffer was prepared in ultrapure water (arium® pro), treated with 0.1 % diethyl pyrocarbonate (D5758, Sigma-Aldrich, Chemie GmbH, Steinheim, Germany) and filter-sterilized (0.22 µM) (Ref: 83.1826.001, Sarstedt AG & Co. KG, Nümbrecht, Germany). The sample was mixed for 3 s. The sample was incubated for 10 min at 42 °C and 850 rpm in the thermomixer to dissolve the pellet. The sample was centrifuged for 30 s. An equal volume (400 µL) of chloroform:isoamyl alcohol (24:1) (v:v) (Carl Roth GmbH & Co. KG; Merck KGaA) was added and the sample was mixed for 3 s. The sample was centrifuged at 20800 x g (Eppendorf 5417 R) at room temperature for 5 min. The upper phase was transferred into a new 1.5 mL reaction tube. The lower phase and interphase were discarded. Another 400 µL of chloroform:isoamyl alcohol (24:1) (v:v) (Carl Roth GmbH & Co. KG; Merck KGaA) were added to the 1.5 mL tube containing the upper phase. The sample was mixed for 3 s and centrifuged at 20800 x g (Eppendorf 5417 R) for 5 min at room temperature °C. The upper phase was transferred into a 1.5 mL reaction tube and if an intermediate phase was visible in the supernatant, the sample was washed again with chloroform:isoamyl alcohol as specified above. Two volumes (800 µL) of cold (- 20 °C) 96% ethanol (Art. No. P075.3, Carl Roth GmbH & Co. KG, Karlsruhe, Germany) were added to the 1.5 mL tube containing the upper phase. The sample was mixed for 3 s. The sample precipitated at -80 °C for 1 h. The sample was centrifuged at

20800 x g (Eppendorf 5417 R) at 4 °C for 20 min. The supernatant was discarded and 500 µL of 70% ethanol (Carl Roth GmbH & Co. KG), which was prepared with molecular grade nuclease-free water (A7398, AppliChem GmbH, Darmstadt, Germany), were added to wash the pellet. The sample was centrifuged at 20800 x g (Eppendorf 5417 R) at room temperature for 10 min. The supernatant was discarded and 80 µL of 70% ethanol (Carl Roth GmbH & Co. KG) were added to wash the pellet again. The sample was centrifuged at 20800 x g (Eppendorf 5417 R) at room temperature for 10 min. The supernatant was discarded, and the sample pellet was placed in a SpeedVac™ Concentrator (Type: 5305, Eppendorf AG, Hamburg, Germany) at 45 °C for 4 min. The pellet was dissolved in 30 µL of molecular grade nuclease-free water (A7398, AppliChem GmbH, Darmstadt, Germany). The sample was mixed for 3 s. To dissolve the pellet, the sample was incubated at 42 °C at 850 rpm for 10 min in the thermomixer. The RNA was aliquoted and stored at - 80 °C. The RNA yield and purity values were measured in a NanoDrop™ 2000/2000c Spectrophotometer (Thermo Fisher Scientific, Wilmington, Delaware USA) and aliquots were used for DNase treatment using the turbo DNA-free kit (Ref: AM1907 from Invitrogen by Thermo Fisher Scientific Baltics UAB, Vilnius, Lithuania) according to the manufacturer's instructions for the "rigorous DNase treatment" protocol.  This approach digests contaminant DNA from the RNA to levels that are below the detection limit by routine PCR. The purified RNA was delivered to Chronix Biomedical GmbH (Goettingen, Germany) for integrity analysis, library preparation and sequencing. The RNA integrity number (RIN) was measured using an Agilent 2100 Bioanalyzer (Agilent Technologies Inc., Santa Clara, California, United States). Twelve samples with RIN value ranging from 6.7 to 7.9 were selected for mRNA library preparation. Libraries were constructed with the NEBNext RNA Ultra II Library Prep Kit for Illumina (New England Biolabs, Ipswich, Massachusetts, United States) from 1 µg of purified RNA according to the manufacturer's instructions. Single-end reads with a length of 75 bp were sequenced on a NextSeq 500 Sequencing System instrument (Illumina, San Diego, CA, USA) with a sequencing depth of 100 million reads per sample, which was applied to capture the fungal mRNA in addition to the beech mRNA in the samples.

**RNAseq bioinformatic processing and data analyses.** Before processing, each sample consisted of around 110 million reads. Processing (trimming, quality filtering and adapter removal) of the raw sequence data was performed using fastp (5). After processing, around 109 million reads per sample remained. The processed RNA reads from our experiment were annotated by mapping against the reference transcriptomes of *Fagus sylvatica* L. and 17 fungi species belonging to the same genus as those detected by metabarcoding. Reference beech sequences and annotations were downloaded from beechgenome.net (13) and reference fungal sequences and annotations were downloaded from the Mycocosm database (mycocosm.jgi.doe.gov) (14). The fungal reference species comprise 13 ectomycorrhizal fungi, one ericoid mycorrhiza, one endophyte, and two saprotrophs: *Amanita muscaria*, *Amanita rubescens*, *Boletus edulis*, *Cenococcum geophilum*, *Cortinarius glaucopus*, *Galerina marginata*, *Laccaria amethystina*, *Laccaria bicolor*, *Lactarius quietus*, *Meliniomyces bicolor*, *Mycena galopus*, *Oidiodendron maius*, *Phialocephala scopiformis*, *Russula ochroleuca*, *Scleroderma citrinum*, *Thelephora terrestris* and *Xerocomus badius.* Although the genus *Boletus* was not detected in beech root apices, the transcriptome of *Boletus edulis* was used as reference to cover the highly abundant Boletales detected in the metabarcoded samples lacking a fine level of taxonomic resolution. The 18 fasta files (fungi + beech) were concatenated to one single file, which was subsequently used to create an index file with bowtie2-build (15). The processed RNA reads were mapped against this index file using bowtie2. On average, 61 % of the RNA reads could be mapped (45 % to beech and 16 % to fungi). After mapping, a combined count table was summarized for beech and for the 17 fungi species from the gene models to which the reads were mapped to. The count table was then split into a separate beech count table and a joint fungi count table. Normalization of the raw count tables and identification of gene models of differentially expressed transcripts of ammonium- or nitrate-treated samples relative to the control was conducted using the DESeq2 package (16), implemented in R (17). Differential expression analyses on the fungi was performed at the metatranscriptome level (i.e, the raw count tables were aggregated by their EuKaryotic Orthologous Groups of protein identifiers (KOGs), thus taxa-specific information for the gene models was dropped). To reduce the number of potential artifacts, gene models having a false discovery rate corrected P $<0.05$ (18) and at least two times more expression ($\log_2$ fold change $<-1.0$ and $>1.0$) were considered statistically significantly differentially expressed (DEGs). To identify associations with the nitrogen metabolism and other metabolic pathways, fungal expressed (i.e., transcribed) genes with assigned Enzyme Commission numbers were mapped to the Kyoto Encyclopedia of Genes and Genomes (KEGG) metabolic pathways against *Laccaria bicolor* in KEGG Mapper (19). Functional enrichment analysis of fungal expressed genes was carried out in g:Profiler (20) against KEGG metabolic pathways with *Aspergillus oryzae* as reference since the model ectomycorrhizal fungus *Laccaria bicolor* was not available. The complete fungal transcriptional database was manually screened for N-related transporters and enzymes using the key words “nitrate transporter,” “nitrate reductase,” “nitrite transporter,” “nitrite reductase,” “ammonium transporter”, “glutamine synthetase,” “glutamate synthase” and “glutamate dehydrogenase.” These terms were searched in the definition lines accompanying the annotations of each of the fungal transcripts: “kogdefline” = definition of the KOG identifiers, “ECnumDef” = definition of EC number, “iprDesc” = description of the InterPro identifiers, and “goName” = description of the Gene Ontology term. For beech, over-representation analysis of biological pathways based on the MapMan bin classification (Ath_AGI_LOCUS_TAIR10_Aug2012) of DEGs was performed using the Classification SuperViewer Tool (21) from the Bio-Analytic Resource for Plant Biology (http://bar.utoronto.ca/). To account for differences in functional profiling programs and their database annotation versions (22), Gene Ontology term enrichment analysis of beech DEGs from the ammonium and nitrate treatment was also performed in g:Profiler (20).

22. Yon Rhee S, Wood V, Dolinski K, Draghici S. 2008. Use and misuse of the gene ontology annotations. Nat Rev Genet. Nature Publishing Group.
